# WDR5 represents a therapeutically exploitable target for cancer stem cells in glioblastoma

**DOI:** 10.1101/2021.09.20.461125

**Authors:** Kelly Mitchell, Sajina Shakya, Sonali Arora, Samuel A. Sprowls, Daniel J. Silver, Christopher M. Goins, Lisa Wallace, Gustavo Roversi, Rachel Schafer, Kristen Kay, Tyler E. Miller, Adam Lauko, John Bassett, Anjali Kashyap, J. D’Amato Kass, Erin E. Mulkearns-Hubert, Sadie Johnson, Joseph Alvarado, Jeremy N. Rich, Patrick J. Paddison, Anoop P. Patel, Shaun R. Stauffer, Christopher G. Hubert, Justin D. Lathia

## Abstract

Glioblastomas (GBMs) are heterogeneous, treatment-resistant tumors that are driven by populations of cancer stem cells (CSCs). Despite their importance for tumor growth, few molecular mechanisms critical for CSC population maintenance have been exploited for therapeutic development. We developed a spatially resolved loss-of-function screen in GBM patient-derived organoids to identify essential epigenetic regulators in the SOX2-enriched, therapy resistant niche and identified WDR5 as indispensable for this population. WDR5 is a component of the WRAD complex, which promotes SET1-family-mediated Lys4 methylation of histone H3, associated with positive regulation of transcription. In GBM CSC models, WDR5 inhibitors blocked WRAD complex assembly and reduced H3K4 trimethylation and expression of genes involved in CSC-relevant oncogenic pathways. H3K4me3 peaks lost with WDR5 inhibitor treatment occurred disproportionally on POU transcription factor motifs, including the POU5F1(OCT4)::SOX2 motif. We incorporated a SOX2/OCT4 motif driven GFP reporter system into our CSC cell models and found that WDR5 inhibitor treatment diminished reporter activity. Further, WDR5 inhibitor treatment altered the stem cell state, disrupting CSC *in vitro* growth and self-renewal as well as *in vivo* tumor growth. These findings highlight the role of WDR5 and the WRAD complex in maintaining the CSC state and provide a rationale for therapeutic development of WDR5 inhibitors for GBM and other advanced cancers.

**Significance:** In this study, we perform an epigenetic-focused functional genomics screen in glioblastoma organoids and identify WDR5 as an essential epigenetic regulator in the SOX2-enriched, therapy resistant cancer stem cell niche.

**Graphical Abstract:** 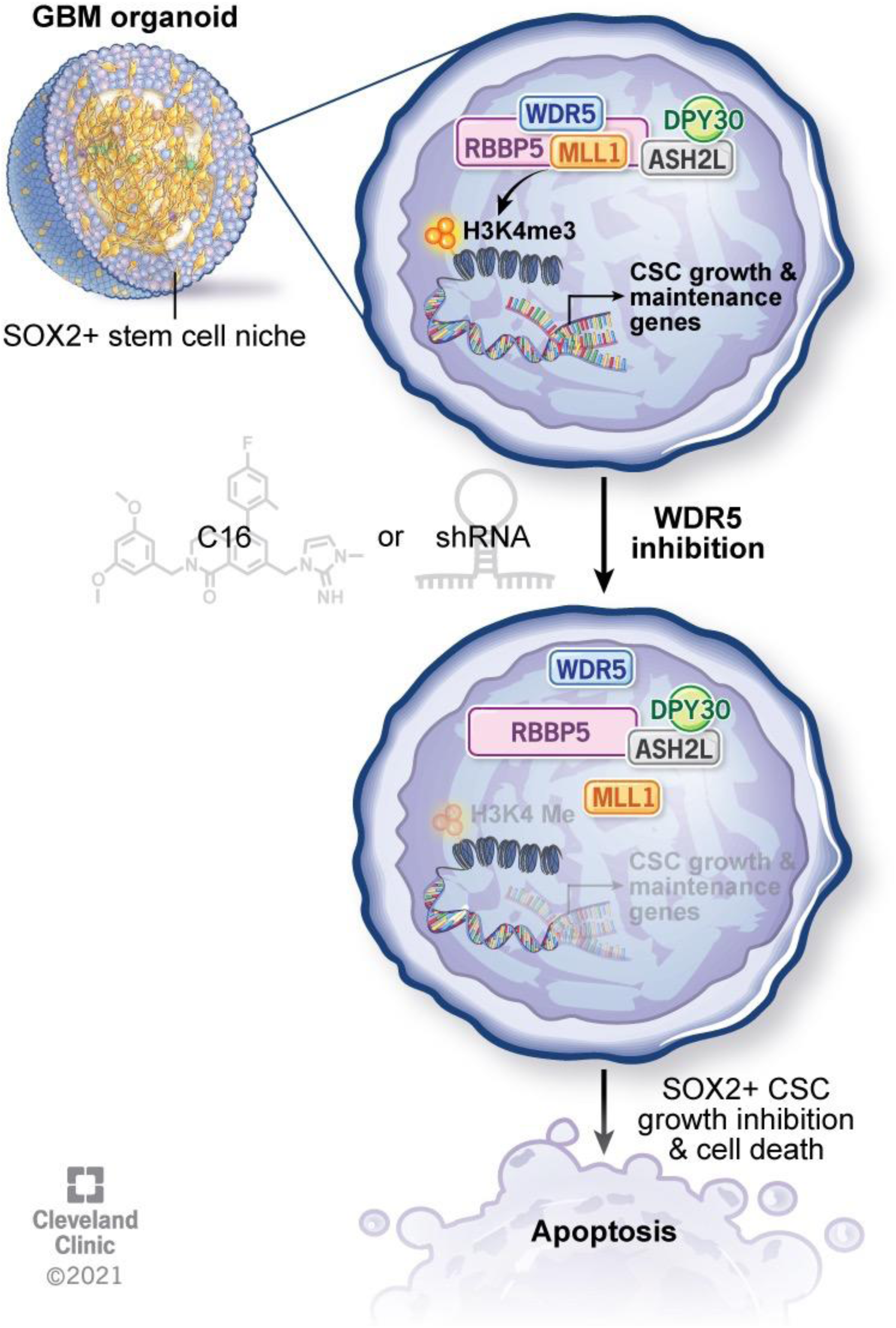

## Introduction

Glioblastoma (GBM) is the most common primary malignant brain tumor and remains highly lethal despite an aggressive multi-modal standard-of-care approach that includes maximal safe surgical resection followed by concomitant radiation and chemotherapy. GBM tumors display profound cellular heterogeneity, with both tumor growth and therapeutic resistance driven by populations of cells with stem-cell like properties that are termed cancer stem cells (CSCs) (reviewed in (Mitchell et al., 2021)). Neurodevelopmental transcription factors such as SOX2, POU3F2, SALL2, and OLIG2 are expressed in subpopulations of GBM tumor cells, are necessary and sufficient for tumor propagation *in vivo*, and cooperate to maintain stem cell-like epigenetic landscapes (Singh et al., 2017; Suva et al., 2014). Such epigenetic regulation controls the access of transcription factors to defined sets of genes and allows for the transition of cells between states (Huang, 2013; Singh *et al*., 2017). Thus, aberrant expression of core transcription factors can coordinate expression of epigenetic regulatory machinery, and together these transcriptional and epigenetic programs promote maintenance of the CSC state in GBM.

Studies profiling DNA methylation and histone modifications in GBM have revealed common transcription factor networks and epigenetic profiles across primary patient tumors (Guilhamon et al., 2021; Hall et al., 2018; Mack et al., 2019; Yoo and Bieda, 2014). Specific DNA methylation, histone methylation, histone acetylation, and chromatin accessibility patterns are predictive of patient response to therapy (Dahlrot et al., 2018; Guilhamon *et al*., 2021; Hall *et al*., 2018; Mack *et al*., 2019; Thon et al., 2013). GBM cell state plasticity in response to external stimuli such as therapeutic pressure is facilitated, at least in part, by chromatin reorganization (Liau et al., 2017).

Accordingly, several epigenetic regulators are elevated in GBM CSCs and are necessary and sufficient for self-renewal. These include retinoblastoma binding protein 5 (RBBP5), which we previously showed to control expression of the core pluripotency transcription factors SOX2, OCT4 and NANOG (Alvarado et al., 2017), mixed lineage leukemia 1 (MLL1) (Gallo et al., 2013; Heddleston et al., 2012), DPY30 (Dixit et al., 2022; Miller et al., 2017), BMI1 (Abdouh et al., 2009), EZH2 (Suva et al., 2009), KDM1A/LSD1 (Suva *et al*., 2014), KDM6B/JMJD3, HELLS, TET3, and TRIM24 (reviewed in (Valor and Hervas-Corpion, 2020)). It is likely that these epigenetic regulators generate permissive chromatin states that then facilitate transcriptional plasticity in tumor cell populations, including maintenance of the CSC state amidst cell-cell interactions, cell-environmental interactions and therapeutic pressures (Flavahan et al., 2017).

3D organoid models of GBM recapitulate a variety of cellular states seen in primary patient tumors, including niches of SOX2+ CSCs, and allow for interaction of cells in various states in an *in vitro* system (Hubert et al., 2016). Primary GBM-derived organoids formed in Matrigel and matured by orbital shaking in serum free, EGF/FGF2 supplemented media show regional heterogeneity with a proliferative, SOX2-enriched outer rim and a hypoxic core harboring quiescent CSCs and non-CSC tumor cells. These organoids recapitulate diffuse and infiltrative properties of human GBM upon xenotransplantation into mice (Hubert *et al*., 2016). To elucidate epigenetic factors responsible for maintenance of the CSC state in GBM in the context of heterogeneous cell populations and CSC niches, we employed an unbiased organoid-based screen targeting epigenetic regulators. Through this screen, we identified WDR5 as a key CSC regulator and validated its functional necessity for CSC self-renewal and tumor initiation.

WDR5 interacts with and facilitates the actions of multiple epigenetic regulator proteins and transcription factors (Guarnaccia and Tansey, 2018). WDR5 is best characterized as a member of the WRAD complex, which also includes the proteins RBBP5, ASH2L, and DPY30, two of which were previously found to be important for the GBM CSC state, including *in vivo* (Alvarado *et al*., 2017; Dixit *et al*., 2022; Miller *et al*., 2017). The WRAD complex interacts with methyltransferases, including MLL1, to facilitate post-translational modifications on histone tails, including histone 3 lysine 4 (H3K4) mono-, di-, and tri-methylation, which are associated with transcriptionally permissive chromatin (Ruthenburg et al., 2006; Santos-Rosa et al., 2002; Wysocka et al., 2005). (Aho et al., 2019b; Lu et al., 2018). Targeting WDR5 provides an alternative approach to directly inhibiting core “stemness” transcription factors, such as the transcriptional master regulator and GBM CSC marker SOX2, that regulate CSCs but are challenging to individually target due to their complex and varied interactions with proteins and DNA (Gangemi et al., 2009). WDR5 is highly conserved and has been demonstrated to regulate developmental differentiation (Ang et al., 2011). It is also functionally important in a variety of cancers, including acute myeloid leukemia (AML) (Aho *et al*., 2019b). Given the importance of WDR5 in transcriptional and epigenetic regulation, here we test the hypothesis that targeting WDR5 could be a means to compromise the ability for GBM CSCs to maintain a favorable epigenetic state.

## Results

### Patient-derived GBM organoid specimens exhibit increased SOX2 expression within the highly proliferative rim region

GBM CSCs reside in defined tumor niches and display complex interactions with their microenvironment and surrounding cell populations (Bayik et al., 2020; Jacob et al., 2020; Lathia et al., 2015; Mitchell *et al*., 2021; Silver et al., 2021; Venkatesh et al., 2015; Zhang et al., 1999). To better model the cellular heterogeneity in GBM and capture the complex dependencies of CSCs, we leveraged an organoid culture system that allows for the simultaneous culture of GBM cells in diverse states, including the CSC state, as previously described (Hubert *et al*., 2016). The outer rim of GBM organoids is highly enriched for functionally self-renewing SOX2+ CSCs compared to the inner hypoxic core, representing two distinct growth zones that support the CSC phenotype to notably different degrees yet arise from a single population of GBM tumor cells (**Fig. 1A-D -** *note: Figs. start on page 56*). To isolate viable GBM populations from these separate zones, we developed a technique to regionally label GBM organoids. This method specifically and reliably labels the outer proliferative niche within each organoid using the fluorescent dye CellTracker Blue CMAC (Shakya et al., 2021). RNA sequencing of spatially isolated GBM cells from dissociated organoids reflects region-specific gene expression profiles from patient GBM tumors, and the SOX2-high organoid outer rim is functionally enriched for stem cell activity (Shakya et al., 2021). This SOX2+ GBM organoid rim is highly proliferative and the SOX2+ cells within this region are resistant to standard of care therapies including radiation therapy (Hubert et al., 2016), the chemotherapy temozolomide, and other clinically relevant therapeutics (Sundar et al., 2021). A wide range of patient-derived GBM organoid specimens demonstrated increased SOX2 expression within the highly proliferative rim region (**Fig. 1E**).

**Figure 1:**
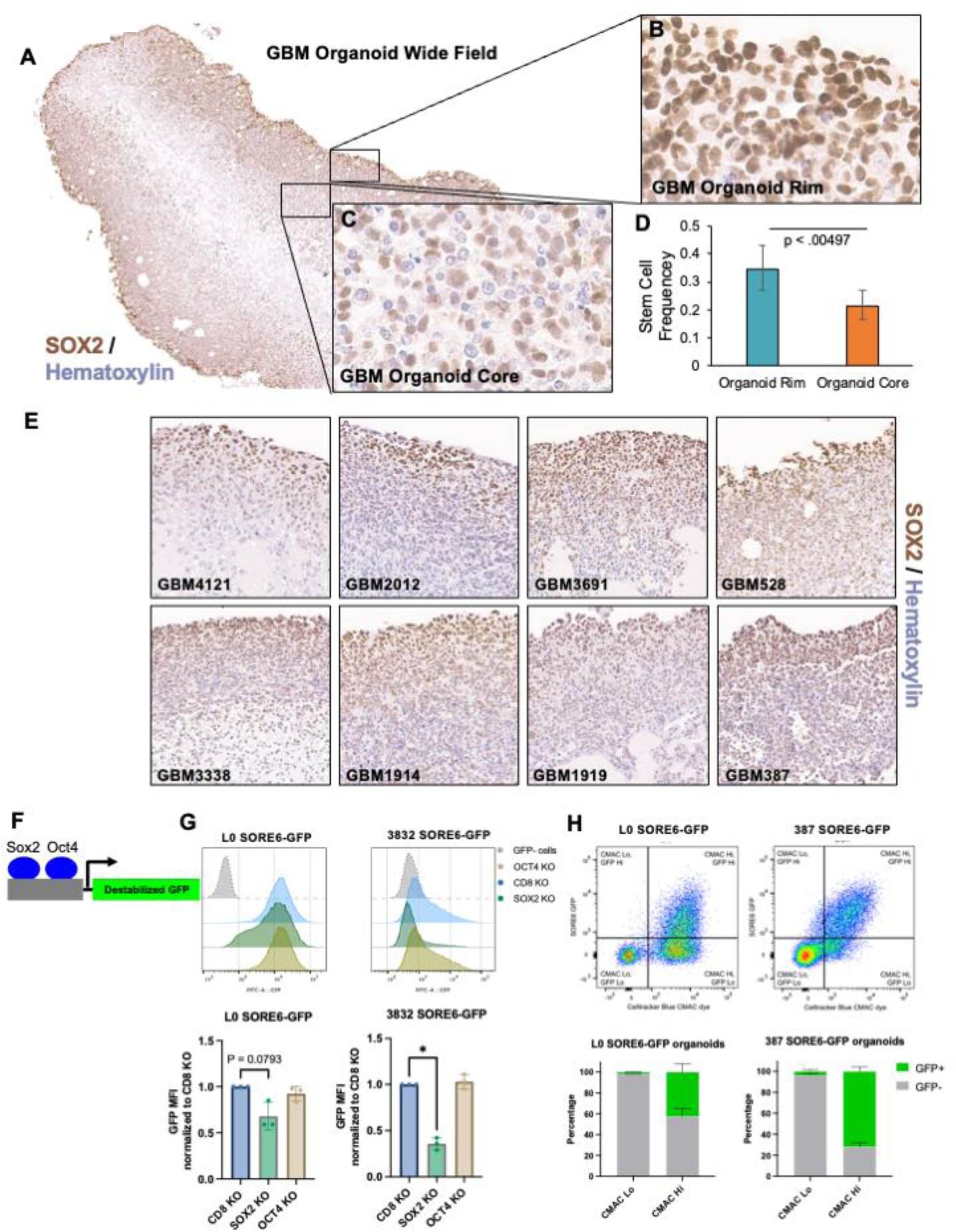
SOX2 is enriched in highly proliferative rim region of patient-derived GBM organoids. **(A-C)** SOX2 IHC shows enrichment of SOX2+ cells in the GBM organoid rim. (**D**) Cells from the organoid rim display a higher frequency of stem cell behavior (sphere formation) by limiting dilution assay. Error bars represent 95% confidence intervals for stem cell frequency. Chi-squared p value for difference in sphere formation is shown. (**E**) SOX2 IHC shows enrichment of SOX2+ cells in the GBM organoid rim of 8 different patient-derived specimens. 20X fields of view for 8 individual specimens are shown. (**F**) Schematic describing the SORE6-GFP lentiviral reporter system. The SOX2 and OCT4 promoter response elements (cloned from the *NANOG* promoter) are fused to a destabilized GFP. (**G**) SOX2 and OCT4 were knocked out via CRISPR:Cas9 in SORE6-GFP transduced GBM CSCs and SORE6-GFP reporter expression was measured by flow cytometry. Bars represent geometric mean fluorescence intensity (MFI) for GFP relative to control, +/-SD; symbols represent biological replicates. P values determined by two tailed, paired t-tests. (**H**) GBM organoids derived from SORE6GFP-transduced GBM CSCs were regionally labeled with the fluorescent dye CellTracker Blue CMAC system, dissociated, and analyzed by flow cytometry to measure GFP expression in the CMAC+ outer organoid niche and CMAC-inner niche. Error bars represent standard deviation of n=3 biological replicates per line.

To specifically measure the abundance of SOX2+ cells in the CMAC-labeled region of organoids, we turned to the lentiviral-based SOX2/OCT4 response element (SORE6) reporter system. In this system, a destabilized copepod-GFP is activated in response to binding to 6 tandem repeats of the OCT4(POU5F1)-SOX2-TCF-NANOG motif (cloned from the NANOG promoter) (**Fig. 1F**) (Tang et al., 2015). Further, the destabilized GFP allows for temporal resolution and real-time monitoring of the activity of these cancer stem cell-relevant transcription factors. To interrogate functionality of the S6GFP system in GBM CSCs, we knocked out SOX2 and OCT4 via CRISPR and monitored the effect on SORE6-GFP reporter expression (**Supp. Fig 1A, B**). In 2 GBM CSC isolates tested, SOX2 KO, but not OCT4 KO reduced GFP (reporter) activity (**Fig. 1G**), suggesting SOX2 is the main driver of the OCT4(POU5F1)-SOX2-TCF-NANOG motif in these cells, and this reporter can be used as a readout of SOX2 activity. To quantify SOX2-expressing and non-expressing cells that result in from CMAC-based labeling of organoid regions, we generated and matured organoids from SORE6GFP-transduced GBM CSCs. Next, we regionally labeled GBM organoids with the fluorescent dye CellTracker Blue CMAC system. We then assessed dissociated organoids by flow cytometry and observed that the CMAC+ outer organoid niche contains the vast majority of GFP+ cells. Only a few percent (about 2%) of cells in the CMAC-group were GFP+, while about half of the CMAC+ cells from the outer region of organoids are GFP+ (**Fig 1H**). This data corroborates our IHC data demonstrating enrichment of SOX2 in the outer niche.

### Spatially resolved organoid screening reveals WDR5 is essential for CSC survival

To elucidate epigenetic regulators responsible for maintenance of the SOX2+, therapy-resistant cellular state, we adapted our methods to enable high-throughput functional screening in 3D organoid culture (**Fig. 2A**). We used a pooled inducible lentiviral shRNA library to target ∼400 epigenetic-modifying genes in GBM CSCs, FACS sorted virus-infected CSCs (mVenus+), seeded CSCs to generate several hundred organoids in parallel, and allowed the organoids to mature for 1 month in spinning bioreactors prior to shRNA induction and outgrowth. We waited 1 month prior to shRNA induction in order to avoid affecting cells prior to stable microenvironment formation (**Supp. Fig. 2A**). We validated viral integration in the entire organoid (mVenus+) and shRNA induction (dsRed+) throughout the entire organoid by microscopy. To isolate separate niche populations, we spatially labeled GBM organoids with CMAC blue as described above and verified proper organoid regional labeling using live confocal imaging (**Supp. Fig. 2B-F**) prior to organoid dissociation and FACS sorting. We sorted on successfully induced mVenus+dsRed+ cells in each region (CMAC blue+ or CMAC blue-), DNA was isolated from sorted populations, tagged with unique molecular barcodes, and deconvolved by high-throughput sequencing of the remaining integrated shRNA libraries as previously described (Miller et al., 2017). Greater than 500-fold representation of library complexity was maintained at every step in the screened populations, and full screens were performed in triplicate. Genes identified by RIGER analysis (Broad Institute) as essential by organoid screening were retrospectively separated into overlapping or niche-specific targets based on prior regional labeling (**Fig. 2B, C, Supp. Fig. 2G**). Our positive control, RPA3 knockdown, is broadly cell lethal and was found to be essential in both cell niche populations. Since cell-lethal knockdowns common to both organoid regions are likely enriched for such universally required genes and therefore less likely to have a therapeutic window upon translation to therapy, we focused on genes uniquely essential in the GBM SOX2+ stem cell niche (**Fig. 2B**). This population included MLL5, a gene previously identified as critical for maintaining CSC self-renewal in GBM (Gallo et al., 2015), underscoring the capability of our screening platform to identify valid and biologically meaningful genes in GBM CSCs.

**Figure 2:**
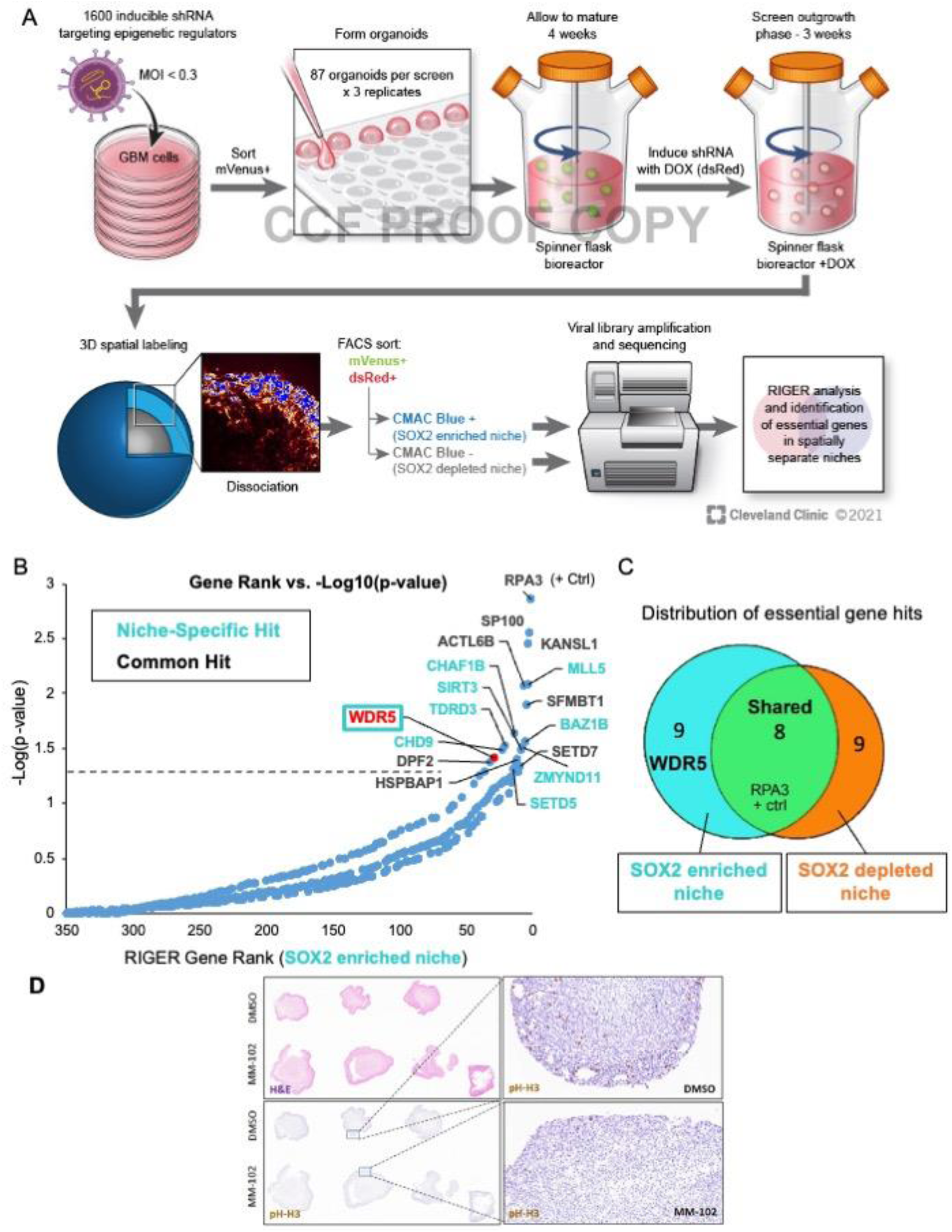
Spatial functional genomics screening defines genes essential in cancer niches. (**A**) GBM CSCs were infected with an inducible shRNA library targeting epigenetic modifiers (mVenus) and grown into organoids. shRNAs were induced with doxycycline (dsRed) and after 3 weeks, organoids were stained with CellTracker CMAC blue dye to label the entire outer rim region. Subsequently, single cells were isolated from the organoids and separated into rim or core populations by FACS, and then DNA was isolated for barcode sequencing and analysis. (**B**) Rank ordered list of genes targeted in the shRNA screen, ranked by depletion of the shRNA as detected by sequencing in the SOX2-enriched niche. Dotted line represents p = 0.05 as determined by RIGER analysis of all hairpin sequences and replicates. Niche-specific hits are color coded, and common hits (including RPA3 positive control) are shown in black. (**C**) Venn diagram of screen hits showing localization in SOX2-enriched or SOX2-depleted niches or common to both regions. (**D**) Organoids derived from GBM CSCs were treated with 63 µM MM-102 for 7 days. Low-power slide scans (left) and 20X fields of view (right) of 3 replicate organoids treated with DMSO or MM-102 showing H&E staining (top left) and phospho-histone H3 staining (right and bottom left) of GBM528 organoids.

Our screen also identified the trithorax protein WD repeat domain 5 (WDR5), a core subunit of the WRAD complex that facilitates activity of human SET1/MLL H3K4 methyltransferase complexes, as an essential gene for growth within the SOX2-enriched niche of GBM organoids. Previous studies demonstrated that WDR5 mediates self-renewal in embryonic stem cells by regulating the OCT4-SOX2-NANOG pluripotency transcription factor network (Ang et al., 2011). While WDR5 has been documented to be expressed in GBM and neuroblastoma and functionally important in high-passage GBM models (Dai et al., 2020; Wang et al., 2020), WDR5 function has not been investigated in the cancer stem cell state in GBM or in GBM organoid culture, and therapeutic inhibition of WDR5 has not been tested in GBM.

Our screen identified KANSL1 as essential in both the SOX2-enriched and SOX2-depleted regions of the organoid. WDR5 is known to bind KANSL1 as an essential part of the non-specific lethal (NSL) histone 4 acetyltransferase complex (Dias et al., 2014). KANSL1 and MLL1 both bind WDR5, mutually exclusively, at its WIN site, an arginine binding cavity. While treatment with a WDR5 WIN site inhibitor, C6 (5 µM for 5 hrs) did not affect KANSL1 binding to flag-tagged WDR5 in 293T lysates (Guarnaccia et al., 2021), WIN site inhibitor, C16 (1 µM for 4 hrs), was shown to inhibit WDR5/MLL1 binding in acute myeloid leukemia cells (Tian et al., 2020). As a tool to inhibit WDR5 without affecting the commonly essential KANSL1, we first treated organoids with a commercially available WDR5 WIN site peptide inhibitor, MM-102, previously shown to inhibit MLL1 histone methyltransferase activity *in vitro* (Karatas et al., 2013). We quantified cell proliferation in each organoid niche region using immunohistochemistry (IHC) for a mitotic marker (phosphorylated histone H3, pHH3) and scanning/tiling microscopy. MM-102 treatment recapitulated the screen results, resulting in reduced pHH3+ cells in the SOX2-enriched niche (**Fig. 2D**).

### WDR5 small molecule inhibition reduces the interaction between WDR5 and WRAD complex members and broadly diminishes histone 3 lysine 4 trimethylation (H3K4me3)

Given our observations that the WDR5 peptidomimetic inhibitor MM-102 reduced proliferation in GBM organoids, we next aimed to determine the molecular effects of WDR5 WIN site inhibition on WRAD complex function. We first sought to validate the interaction between WDR5 and members of the WRAD complex in these models. By immunoprecipitation of either RBBP5 or WDR5 and immunoblotting, we detected the association of WDR5 with RBBP5 and the other WRAD complex members in CSCs (**Fig. 3A, Supp. Fig. 3A**). Considering that MLL1 is specifically dependent on WDR5 for its methyltransferase activity (Alicea-Velazquez et al., 2016; Cao et al., 2014; Dou et al., 2006; Shinsky et al., 2015) and given the previously described role of MLL1 in GBM CSCs (Gallo *et al*., 2013; Heddleston *et al*., 2012), we additionally immunoblotted for MLL1 and found that it was bound to WDR5 and RBBP5 (**Fig. 3A, Supp. Fig. 3A**). To assess WDR5 WIN site inhibition in the context of this complex, we turned to compound 16, a recently disclosed small molecule WDR5 WIN-site inhibitor (Tian *et al*., 2020) (**Fig. 3B**), for brevity referred to as C16 hereafter. We chose C16 as it was synthesized in a series of compounds that have improved on-target potency and drug-like properties compared to previously described WDR5 WIN site inhibitors. C16 was the most potent in this series of compounds in its picomolar binding affinity for WDR5 and low nanomolar inhibition of MLL1 histone methyltransferase activity. In addition, an X-ray co-crystal structure of C16 bound to WDR5 has been solved (Tian et al., 2020). We synthesized C16 using published protocols (Tian et al., 2020). Using recombinantly expressed and purified WDR5, the C16 small molecule inhibitor robustly displaced a fluorescently labeled MLL1-derived peptide using the time-resolved fluorescence energy transfer (TR-FRET) assay (**Supp. Fig. 3B**). To gain further insight into the effects of C16 in GBM CSCs, we performed co-immunoprecipitation studies in 3 GBM CSC models after C16 treatment with a focus on MLL1 and core WRAD complex members (schematic in **Fig. 3B**). Unlike for other SET1 family methyltransferases (SET1A, SET1B, MLL2-MLL4), WDR5 is indispensable for MLL1’s methyltransferase activity (Li et al., 2016; Zhang et al., 2018). In addition, only MLL1 and SET1A were shown to be specifically reliant on the WDR5 WIN site interaction for methyltransferase activity (Alicea-Velazquez *et al*., 2016; Li *et al*., 2016; Patel et al., 2008). As expected (Tian *et al*., 2020), MLL1 was displaced from WDR5 and RBBP5 upon C16 treatment. However, in contrast to what was found in AML cells (Tian et al., 2020), the interaction between WDR5 and RBBP5 was also reduced (**Fig. 3C,D, Supp. Fig. 3C,D**). C16 targets the WDR5/MLL1 interaction site, which is distinct from the WDR5/RBBP5 interaction site, yet the RBBP5 interaction appears to be allosterically affected in our GBM CSC models. Previous studies showed that WDR5 inhibitor OICR-9429 reduced the amount of endogenous RBBP5 that co-immunoprecipitated with FLAG-tagged (exogeneous) WDR5 in HEK293 cells (Grebien et al., 2015), but similar results were not seen with the WDR5 WIN site inhibitor C6 in HEK293-WDR5-FLAG cells (Guarnaccia *et al*., 2021) or with C16 in AML cells (Tian et al., 2020). Thus, WDR5/WIN site inhibition may have different effects in different tumor contexts and effects likely depend on the specific inhibitor.

**Figure 3:**
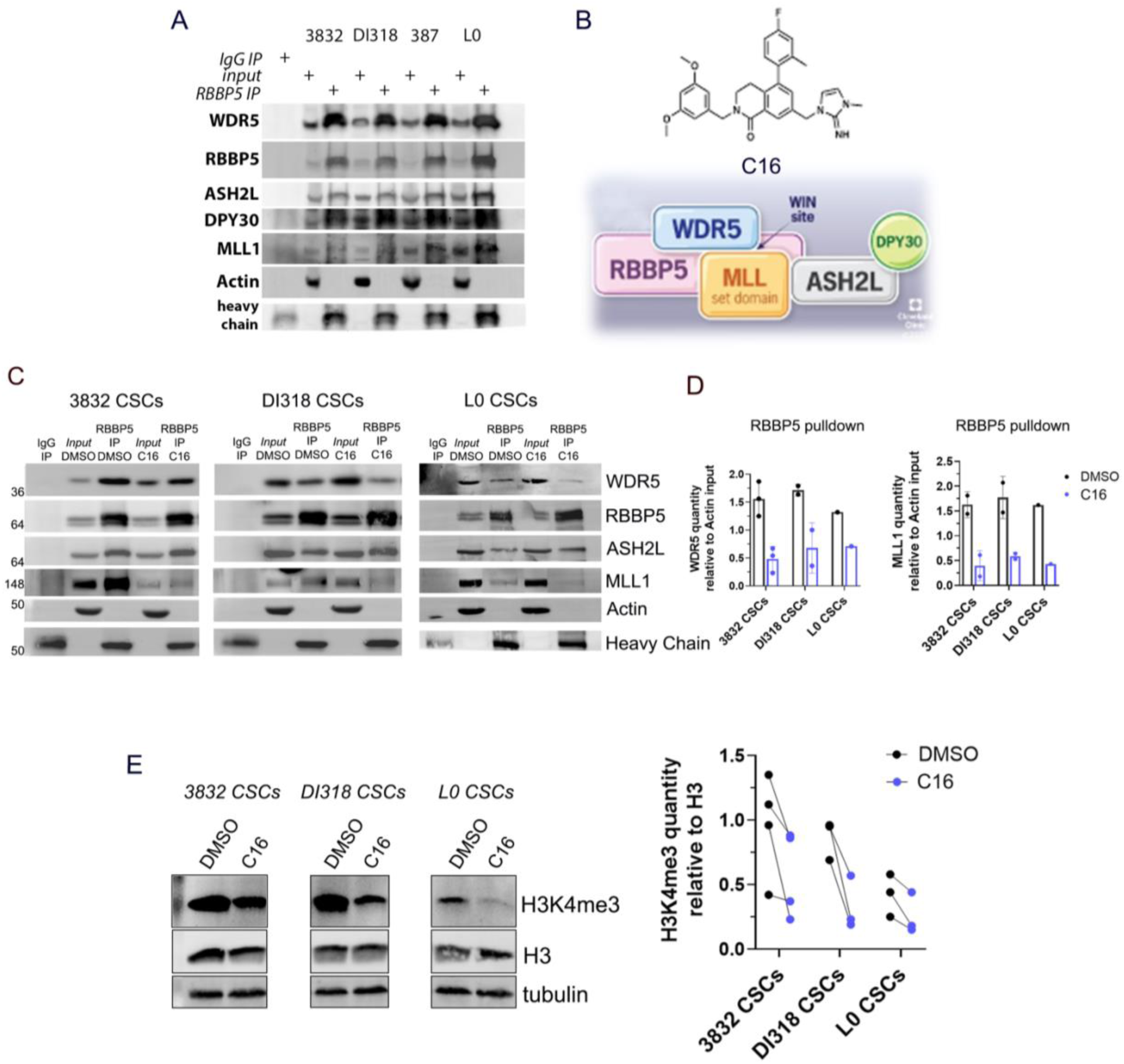
WDR5 inhibition reduces the interaction between WDR5 and WRAD complex members and diminishes the H3K4me3 mark. (**A**) Immunoprecipitation of RBBP5 in 4 GBM CSC models. Immunoblotting was performed for WRAD complex members and the WRAD-associated methyltransferase MLL1. Inputs are 10%. (**B**) Structure of small molecule WDR5 WIN site inhibitor C16 (top) and model of the WRAD complex indicating points of protein-protein interactions, based on structures solved in (Xue et al., 2019) (bottom). (**C**) Immunoprecipitation of RBBP5 after C16 inhibitor treatment (5 µM, 24 hrs) in 3 GBM CSC models. Immunoblotting was performed for WRAD complex members and the WRAD-associated methyltransferase MLL1. Inputs are 10%. Representative experiments are shown. (**D**) Related to 2B; quantification of WDR5 (left) and MLL1 (right) immunoprecipitated by RBBP5, relative to actin quantity from each sample’s input. Circles represent biological replicates. (**E**) Left: Western blots showing histone 3 lysine 4 trimethylation (H3K4me3) levels in whole cell lysates after C16 treatment (5 µM, 72 hrs) in 3 CSC models. Representative experiments are shown. Right: Quantification of H3K4me3, relative to H3 quantity after C16 treatment (5 µM, 72 hrs) in 3 CSC models. Circles represent biological replicates; lines connect DMSO- and C16-treated specimens from the same experiment.

Due to our observation of MLL1 dissociation from RBBP5/WDR5, we next aimed to assess whether MLL1 function is affected by C16 in our GBM CSCs. We first monitored for global changes in H3K4me3. After treatment with C16 for 72 hrs (to allow sufficient time for histone modification changes to occur) we observed a global reduction in H3K4me3 by western blot (**Fig. 3E**). These data corroborate previous studies that showed WDR5 depletion or WDR5 inhibition led to genome-wide reduction in H3K4me3 (Benayoun et al., 2014; Siladi et al., 2022; Zhang et al., 2018). Based on these changes, we utilized the Cleavage Under Targets and Tagmentation (CUT&Tag) approach to identify changes in H3K4me3 localization and abundance after C16 treatment. CUT&Tag allows for efficient profiling of chromatin modifications with low background signal and reduced sequencing depth required to identify histone post-translation modification profiles (Janssens et al., 2018; Kaya-Okur et al., 2020; Kaya-Okur et al., 2019; Lee et al., 2005; Wu et al., 2021). We treated DI318 CSCs, a newly derived CSC model from a GBM patient specimen, with 3 μM C16 or vehicle for 72 hrs and subjected these cells to CUT&Tag for H3K4me3. After performing quality control on the sequencing data, genome alignment, filtering and conversion as previously described (Henikoff et al., 2020), we utilized MACS2 for peak calling and found that replicates from each group clustered closely together based on similarities of peak sequences (**Fig. 4A**). In DMSO treated cells, we identified an average of 21,224 H3K4me3 peaks across replicates, while in C16 treated cells, we identified an average of 19,502 peaks across replicates. We first sought to identify peaks that disappear or appear following C16 treatment and are therefore unique to each group. After excluding peaks less than 50 bp, we identified 1110 peaks unique to the DMSO group (corresponding to 995 genes) and 783 peaks unique to the C16 group (corresponding to 758 genes) (**Fig. 4B, Supp. Table 1**). The peaks lost with C16 treatment were generally larger than the peaks gained with C16 treatment (**Supp. Fig. 4A**). H3K4me3 is enriched immediately downstream of TSSs (Guenther et al., 2005), thus, as expected, the majority (72%) of peaks lost with C16 treatment were contained within 1 kb of the transcription start sites (TSSs). Peaks in exons and in distal intergenic regions made up 20% of peaks lost (**Fig. 4C**). Peaks unique to the DMSO group were enriched for biological processes including neurogenesis and cell projection. Meanwhile, peaks unique to the C16 group were enriched for catabolic processes, tube formation, intracellular protein transport and RNA metabolic processes (**Fig. 4D**). The cellular processes enriched among peaks unique to the C16 group suggest there are compensatory metabolic and developmental pathways being upregulated in response to WDR5 inhibition.

**Figure 4:**
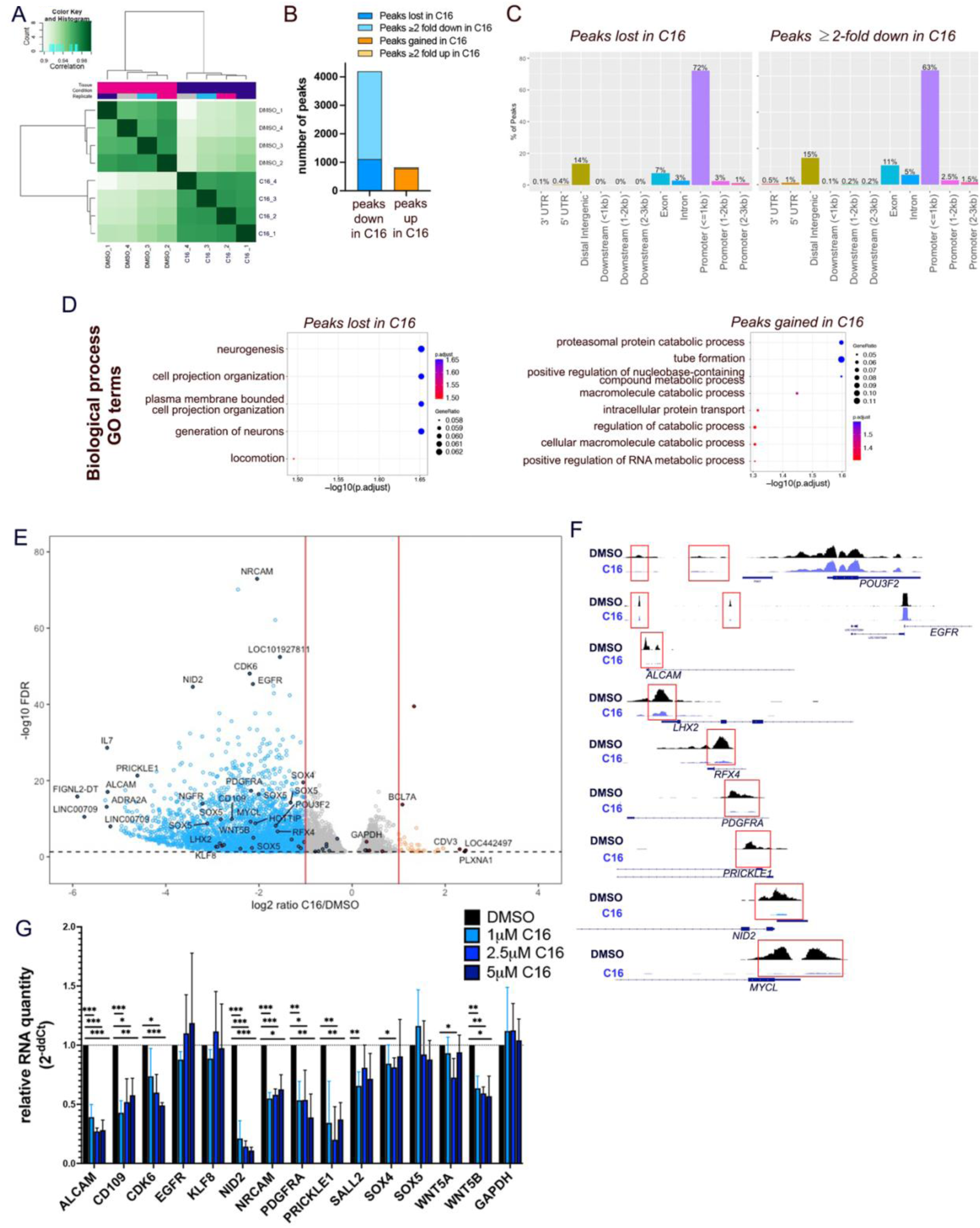
WDR5 inhibition leads to H3K4me3 loss at essential CSC genes and preferentially at POU-domain DNA binding motifs. **(A)** Hierarchically clustered correlation matrix to evaluate the relationship between CUT&Tag replicates. Pearson correlation of the log2-transformed values of read counts in each 500 bp bin between replicates. **(B)** Number of CUT&Tag peaks lost, gained, decreased or increased (log2FC≤-1 or ≥1) after C16 treatment of DI318 cells (3μM for 72 hrs). (**C**) Bar plot showing distribution of peaks in gene regions that were identified as unique to the DMSO group or ≥2-fold downregulated in the C16 treatment group. **(D)** MSigDB gene set annotations enriched among CUT&Tag peaks lost in C16 treatment group (unique to DMSO) and gained in C16 treatment group (unique to C16). **(E)** Volcano plot of differential H3K4me3 peaks detected by CUT&Tag in DI318 CSCs. Blue dots represent H3K4me3 peaks reduced (log2FC≤-1) with C16 treatment and orange circles are H3K4me3 peaks increased (log2FC≥1) with C16 treatment. For some genes, multiple peaks (circles) may map to the same gene. (**F**) Representative H3K4me3 peaks from CUT&Tag at indicated genes. (**G**) qPCR for specified genes on DI318 CSCs treated with indicated doses of C16 for 72 hrs. Bars represent mean expression of n=3 biological replicates, normalized to ACTB levels by ddCt method, +/-SD.

Since the majority of H3K4me3 peaks were still present after C16 treatment, we next sought to identify consensus peaks and determine whether there was differential enrichment of these peaks between the DMSO and C16 groups (**Supp. Table 2**). For this differential analysis, we utilized DiffBind (Ross-Innes et al., 2012) and DESeq2 (Love et al., 2014). 77% of consensus peaks were contained within 1 kb of the TSSs (**Supp. Fig. 4B**). Of all consensus peaks with FDR<0.05 (7,728 peaks), 3,085 were diminished at least 2-fold in the C16 group compared to DMSO group (**Fig. 4B,E**). These 3,085 peaks corresponded to 2,454 genes, 599 of which were the same genes that lost a peak with C16 group (**Supp. Fig. 4C**). Only 39 peaks were increased 2-fold or greater in the C16 treatment group, corresponding to 37 genes, 10 of which were the same genes that gained a peak in the C16 treatment group (**Supp. Fig. 4D**). Genes with peaks “down” in the C16 group were mostly unique from those with peaks “up” in this group compared to DMSO, as only 49 genes both lost and gained a peak in the C16 group compared to the DMSO group and only 3 genes displayed both a 2-fold downregulated peak and a 2-fold upregulated peak in the C16 group compared to the DMSO group. As with peaks lost with C16 treatment, the majority (63%) of ≥2-fold diminished peaks were contained within 1 kb of TSSs, followed by distal intergenic regions (15%) and exons (11%) **(Fig. 4C**). Loss of H3K4me3 in C16 treated CSCs occurred on genes with previously described roles in GBM, such as ALCAM, CD109, EGFR, KLF8, NRCAM, PDGFRA and SOX4 (Filppu et al., 2021; Furnari et al., 2015; Jackson et al., 2006; Kijima et al., 2012; Schnell et al., 2012; Sehgal et al., 1999; Zhang et al., 2014), and CSC-specific master transcription factors LHX2, MYCL, RFX4, CITED1, HEY2, SOX5, POU3F2 and SALL2 (**Fig. 4E, F**), of which the latter two are essential for GBM propagation (Suva *et al*., 2014). H3K4me3 peaks diminished after C16 treatment were enriched for brain developmental and differentiation pathways, synaptic signaling, small GTPase signal transduction, and oncogenic signatures including KRAS, E2F3 (involved in G1/S transition), EGFR and LEF1, among others. WNT/Frizzled binding and the WNT signaling pathway were also enriched among genes with diminished H3K4me3 after C16 treatment (**Supp. Fig. 4E-H)**. Interestingly, WDR5 has been previously implicated in regulating β-catenin transcription via interaction with the long non-coding RNA HOTTIP (Liu et al., 2020), whose corresponding gene also displayed loss of H3K4me3 in C16 treated CSCs (**Fig. 4E**, **Supp. Table 2**). We measured expression of several of these targets at the mRNA level and found that some, but not all, CSC-relevant genes with H3K4me3 loss also had reduced expression with C16 treatment (**Fig. 4H, Supp. Fig. 4I**). For some CSC-specific master transcription factors, such as EGFR, while H3K4me3 peaks >1kb upstream of TSSs were reduced with C16 treatment (**e.g. Fig. 4F**), RNA levels were not significantly reduced (**Fig. 4G**). It is likely that due to the stringent regulation of these key transcription factors, multiple chromatin modification changes are needed to significantly alter gene expression.

### CUT&Tag reveals loss of H3K4me3 at specific POU-domain DNA binding motifs

To ask mechanistically which gene regulatory programs are most affected by WDR5 perturbation in CSCs, we used HOMER analysis within the H3K4me3 peaks that were reduced ≥2-fold in the C16 treatment group to identify enriched DNA binding motifs. The most significantly changed motifs (q-value <0.02) correspond to proteins known to play key roles in glioma maintenance including OCT4 (POU5F1), SOX2, POU3F3 (BRN1), CTCF, and REST, and were strongly enriched (H.Geo. p(X≥7) < 5.7×10^−7^) for members of the POU-domain transcription factor family, known to maintain pluripotency in embryonic stem cells and self-renewal in multiple normal and cancer stem cell types (**Fig. 5A**). A previous study showed that in embryonic stem cells, WDR5 and the POU transcription factor OCT4 interact, and DNA specificity conferred by OCT4 directs WDR5 to specific loci, namely those driving self-renewal (Ang *et al*., 2011). As it is currently unknown how WDR5 is specified to histone targets in GBM, our data provide insight into factors that may cooperate with WDR5 in promoting activation of target gene programs.

**Figure 5:**
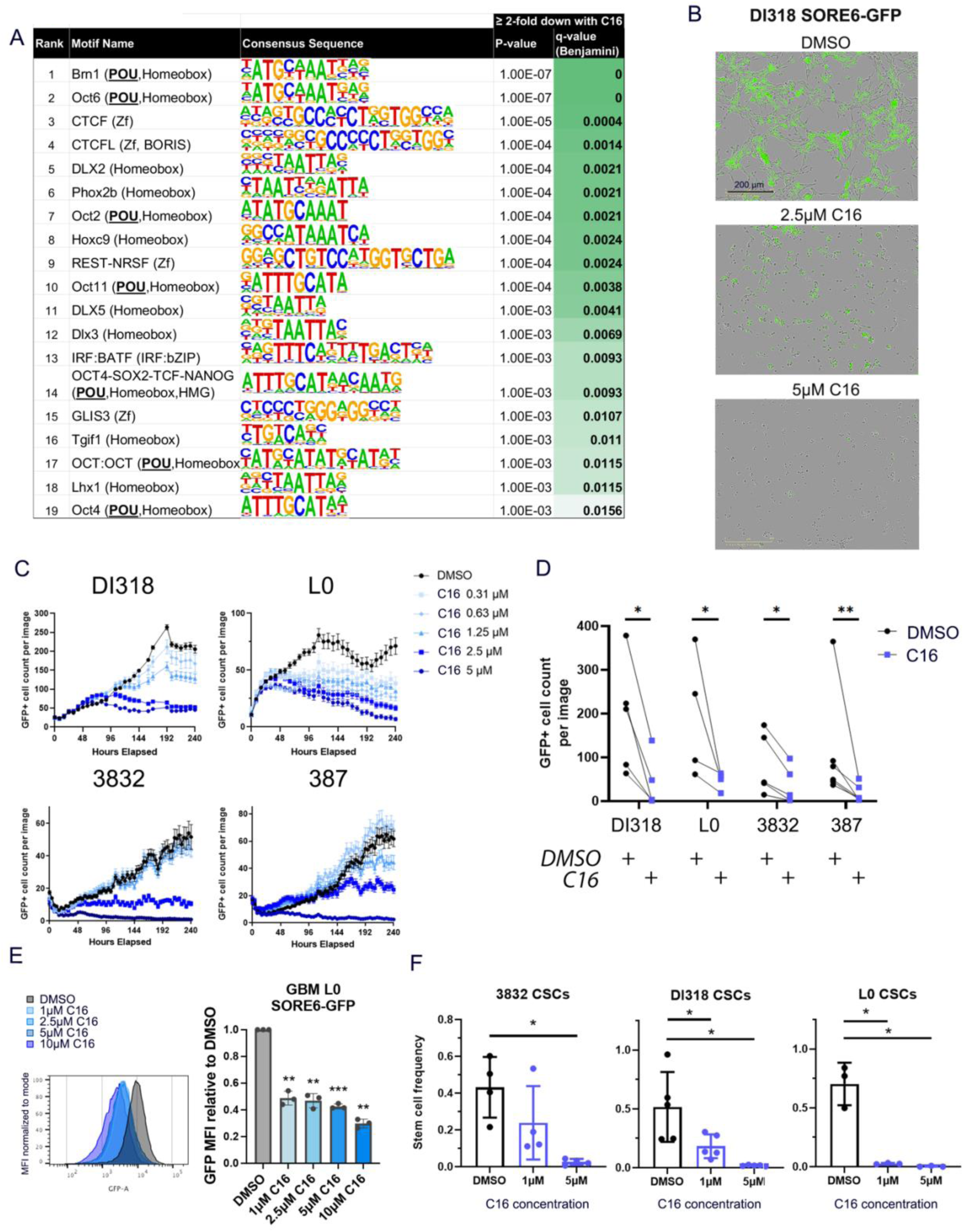
WDR5 inhibition diminishes SORE6 reporter activity and GBM CSC self-renewal. (**A**) Top DNA binding motifs enriched within H3K4me3 peaks reduced (log2FC≤-1) with C16 treatment. The enrichment p-values were computed using the HOMER motif analysis toolset with all significant peaks from the DiffBind analysis as background. (**B**-**D**) GBM CSC models transduced with the SORE6-GFP reporter were treated with the C16 WDR5 inhibitor. (**B**) Representative images Day 7 post treatment are shown. (**C**) GFP+ cell numbers were quantified over 10 days using IncuCyte live cell imaging. One representative time course experiment is shown for each CSC model. Average GFP+ cell count per image is plotted at each time point, +/-SEM per image. Multiple images were taken per well with n=3 technical replicates (wells). (**D**) GFP+ cell numbers at Day 7 after treatment with 5 µM C16. Each line represents a biological replicate. p values determined by two-tailed, paired t-tests. (**E**) GFP intensity of live L0 SORE6GFP cells after C16 treatment for 3 days. P values were determined by two-tailed, paired t-tests. **p<0.01, ***p<0.001. (**F**) *In vitro* limiting-dilution analysis was performed on CSCs in the presence of C16. Bars represent mean sphere formation frequency, +/-SD; symbols represent biological replicates. A table with mean sphere formation frequency per group is shown in Supplemental Figure 5E.

Among the few DNA motifs that significantly lost H3K4me3 after WDR5 inhibition was the OCT4-SOX2-TCF-NANOG (POU, Homeobox, HMG) motif (p <0.001, q <0.0093). In light of this result, and our original identification of WDR5 as critical in the SOX2-enriched GBM niche, we were thus interested in testing whether WDR5 inhibition affects sites regulated by the core stem cell transcription factors SOX2 and OCT4. We first validated the expression of SOX2 in our CSC-enriching culture conditions. We detected SOX2 expression in our CSC models in CSC-enriching culture conditions, and this expression was reduced in serum conditions. SOX2 expression was also detected, albeit at lower levels, in immortalized neural stem cell lines (**Supp. Fig. 5A**). To mechanistically test this motif as a WDR5-regulated sequence, we utilized the SORE6 reporter system (**Fig. 1F**). While this system has been utilized as a tool in a variety of tumors (Koshkin et al., 2021; Li et al., 2021; Menendez et al., 2020; Padua et al., 2020; Pudelko et al., 2017; Tang *et al*., 2015; Vaddi et al., 2019), it has not yet been employed to inform the GBM CSC phenotype. Live cell imaging of SORE6-reporter-transduced CSCs revealed that C16 treatment decreased the number of GFP+ cells over time, indicating WDR5 inhibition diminished reporter activity (**Fig. 5B-D, Supp. Fig. 5B**). SORE6-GFP^high^ cells gave rise to both GFP^high^ and GFP^low^ cells, with GFP^high^ cells having a higher self-renewal frequency (**Supp. Fig. 5C**), as expected for cells with high SOX2/OCT4 activity. To validate the loss of GFP intensity is a result of inhibited transcription but not cell death, mean GFP intensity of live cells was compared between control and C16-treated GBM CSCs expressing the SORE6-GFP reporter. GFP intensity of live SORE6GFP cells was reduced after C16 treatment (3 day treatment) (**Fig. 5E**).

To further interrogate the CSC state in response to WDR5 inhibition, we performed limiting dilution sphere formation assays and found C16 treatment reduced CSC self-renewal in a concentration-dependent manner (**Fig. 5F, Supp. Fig. 5D,E**). Importantly, we observed loss of proliferation specifically in SOX2+ cells in organoids with C16 treatment (**Supp. Fig. 5F**). Taken together, these data show that WDR5 inhibition turns off SOX2/OCT4-regulated loci and provide proof of concept that targeting of the WDR5 suppresses the CSC phenotype by disrupting epigenetic programs that maintain the CSC transcriptional state.

### Reduction of CSC growth and viability via WDR5 inhibition

Given our initial functional and mechanistic assessments of the WDR5 inhibitors MM-102 and C16, we aimed to further assess the effects of these and other WDR5 inhibitors in CSC-enriched PDX cultures. We turned to a series of patient-derived GBM xenograft (PDX) models using conventional CSC-enriching culture conditions to more efficiently expand and validate the function of WDR5 in CSCs. We observed a reduction in CSC number and proliferation upon treatment with C16 and MM-102 in a dose-dependent manner (**Fig. 6A, Supp. Fig. 6A,B**). For C16, half-maximal inhibitory concentrations (IC_50_) values ranged from 0.4-6.6 µM across CSCs from >8 PDX models. Meanwhile, the IC_50_ values for MM-102 ranged from 20-40 µM (**Table 1**), which we predict would pose a challenge for eventual clinical translation. As a large peptidomimetic, MM-102 is limited to an *in vitro* setting and clearly lacks the properties needed (e.g., passive permeability, CNS penetration, metabolic stability) to fully study the impact of WDR5 WIN-site inhibition in the context of GBM. Thus we also obtained two previously described small molecule WDR5 inhibitors, piribedil and OICR-9429. These have been tested in multiple cancers, including AML, neuroblastoma and pancreatic ductal adenocarcinoma (Aho et al., 2019a; Sun et al., 2015; Tian *et al*., 2020; Zhang *et al*., 2018), but they have not been assessed in GBM or CSCs. These two compounds displayed IC_50_s similar to those observed with MM-102 (**Table 1**). With WDR5 binding affinity confirmed for C16 (**Supp Fig. 3B**) and since GBM CSC models were more sensitive to C16 compared with the other WDR5 inhibitors tested (**Table 1**), we moved forward with further testing of C16.

**Table 1:**
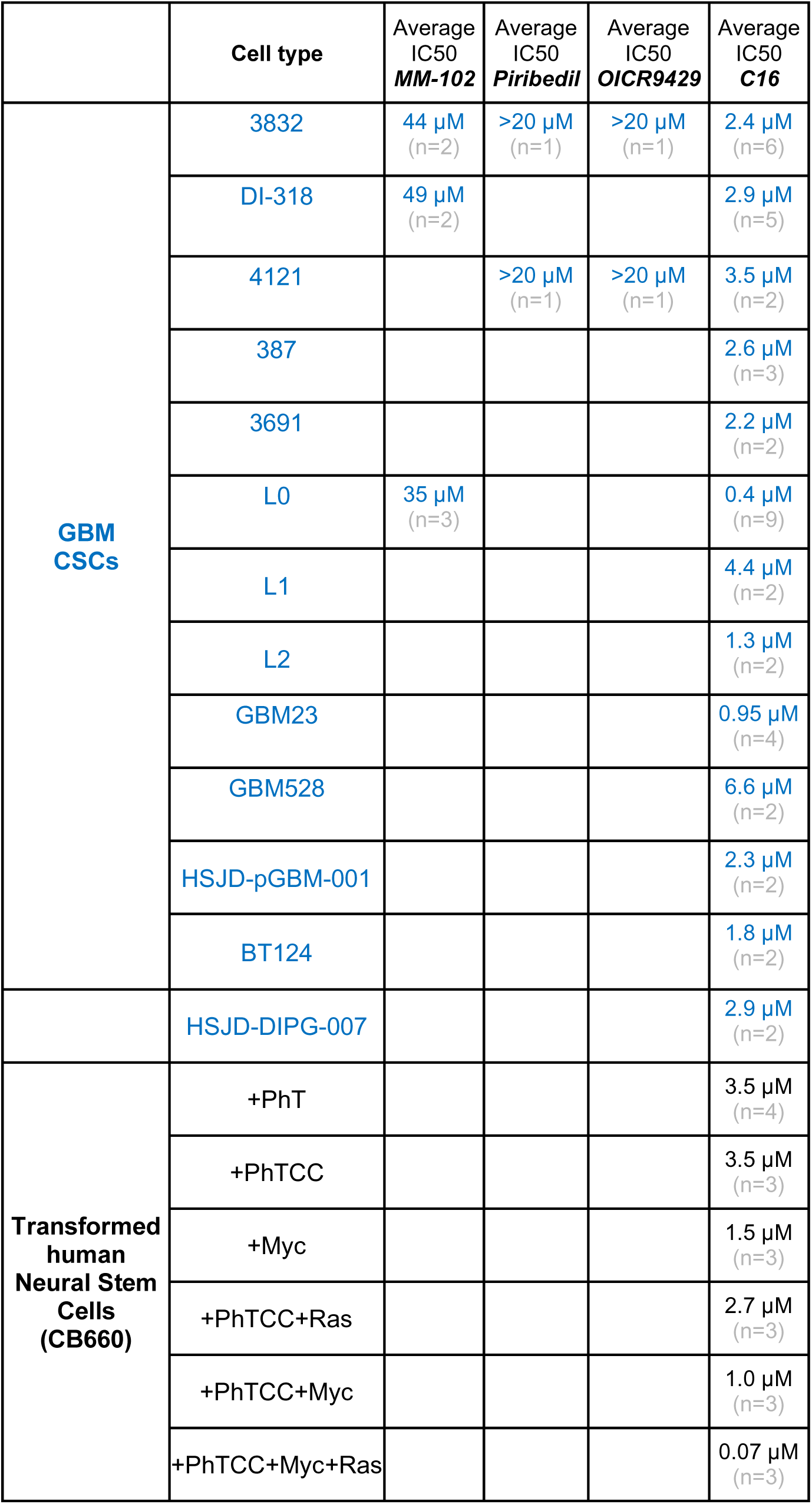
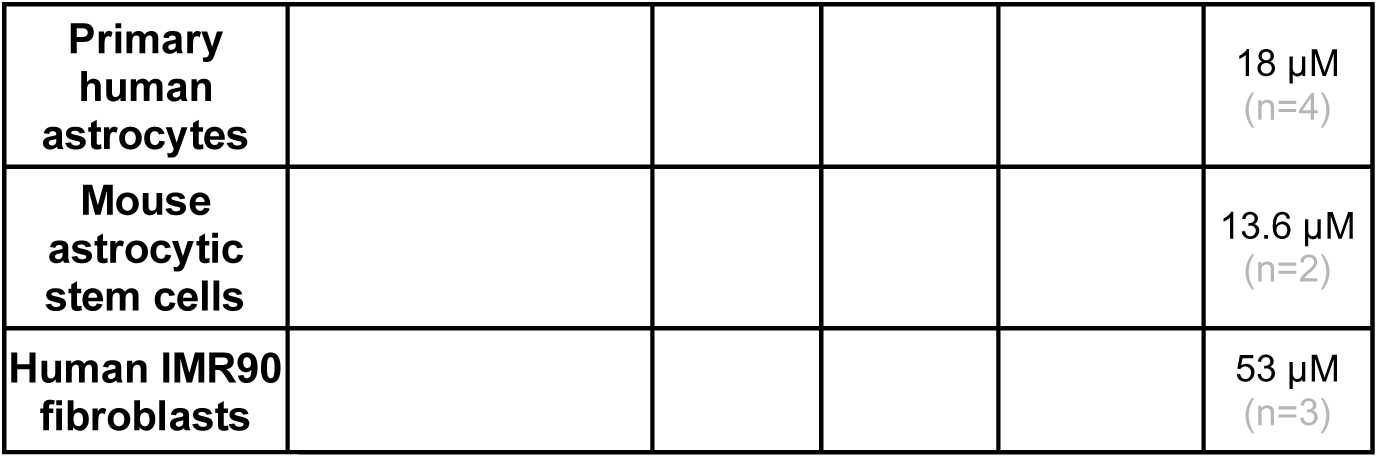
IC50 values of WDR5 inhibitors on GBM CSC models and control cell types. Cells were treated with a range of concentrations of the peptide WDR5 inhibitor MM-102 and small molecule WDR5 inhibitors Piribedil, OICR9429 and C16. After 7 days, viable cell counts were measured by CellTiter Glo viability assay. Values represent mean IC_50_ values (relative to DMSO-treated cells) across multiple experiments; number of independent replicates is indicated in the table.

**Figure 6:**
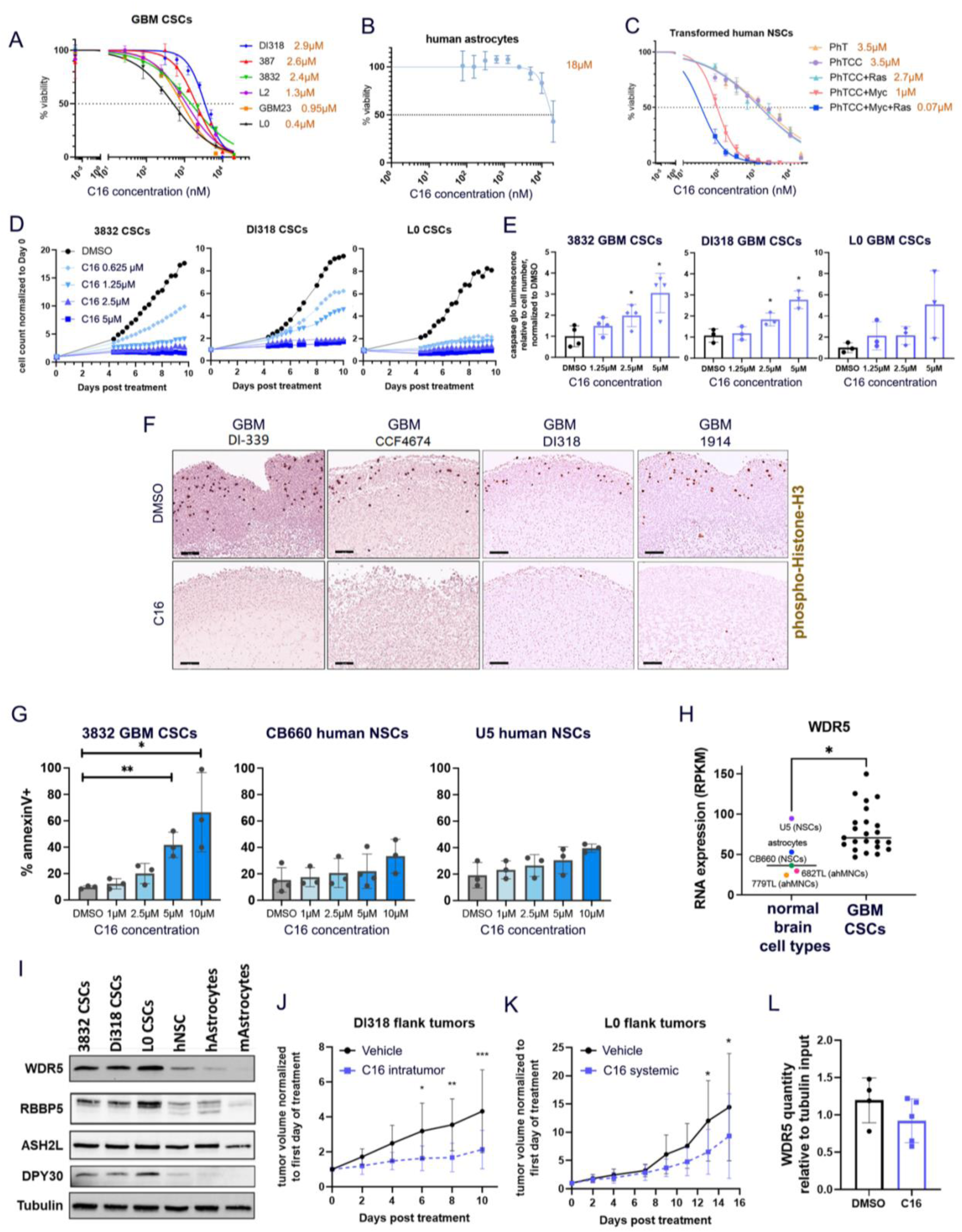
A WDR5 small molecule inhibitor reduces GBM CSC growth and viability. (**A-C**) GBM CSCs (**A**) and human astrocytes (**B**) and CB660 transformed human neural stem cells (**C**) were treated with a range of concentrations of C16, a small molecule inhibitor of WDR5. After 7 days, viable cell counts were measured by CellTiter Glo viability assay. Values represent mean luminescence values normalized to DMSO-treated cells. One representative curve per cell model is shown. Average IC50 values over multiple independent replicates is shown in orange, number of replicates is shown in Table 1. (**D**) Proliferation of C16-treated CSCs over 10 days, determined by IncuCyte live cell imaging. Values represent mean fold change in cell count relative to Day 0, +/- SD, n=3 technical replicates; one representative experiment is shown per CSC model. (**E**) GBM CSCs were treated with a range of concentrations of C16 and subjected to Caspase 3/7 Glo luminescence assay after 4 days to measure caspase 3/7 activity. Bars represent fold change in caspase 3/7 activity per cell relative to the average for DMSO-treated cells, +/- SD; circles represent biological replicates. p values determined by one way ANOVA and post-hoc Dunnett’s multiple comparisons test. (**F**) IHC staining of mitotic marker phospho-histone H3 (pHH3) in 4 independent C16-treated GBM organoids (10 µM for 7 days) at 10X magnification. (**G**) human GBM CSCs or NSCs were treated with WDR5 inhibitor C16 for 4 days and assessed for apoptosis by annexinV and DAPI staining by flow cytometry. Error bars represent +/- SD; circles represent biological replicates. p values determined by one way ANOVA and post-hoc Dunnett’s multiple comparisons test. (**H**) WDR5 expression from RNA sequencing of a panel of normal brain cell types and GBM lines (from (Toledo et al., 2015). Each point represents average expression from multiple sequencing replicates. (**I**) WRAD complex member expression in 3 CSC models, transformed human neural stem cells (hNSCs), human (h) and mouse (m) astrocytes. (**J**) A total of 500,000 DI318 CSCs were implanted into the flanks of mice, and once tumors developed, 3 mg/kg C16 was injected into the tumors daily. Tumor volume over time normalized to tumor size at Day 0 (left) is shown. p values determined by two-tailed, unpaired t test comparing means per group at each time point. n=10 per group. (**K**) A total of 500,000 L0 CSCs were implanted into the flanks of mice, and once tumors developed, 10 mg/kg C16 was injected intraperitoneally daily. Tumor volume over time normalized to tumor size at Day 0 (left) is shown. p values determined by two-tailed, unpaired t test comparing means per group at each time point. n=9 for DMSO group, n=10 for C16 group. (**L**) Immunoprecipitation of RBBP5 and immunoblot for WDR5 was performed on flank tumor lysates isolated from systemic vehicle and C16 treated mice in 3H. Symbols represent individual mice. WDR5 quantity in RBBP5 pulldown lanes relative to tubulin quantity in input (10%) lanes is plotted.

In light of the previously described role of the WRAD complex in embryonic stem cells (Ang *et al*., 2011; Lim et al., 2009), we aimed to test the effect of C16 in normal brain cell types. We tested C16 on astrocytes and fibroblasts *in vitro* to assess toxicity on relevant normal cell populations. IC50 estimates of normal cell types treated with WDR5 inhibitor C16 for 7 days revealed that human astrocytes had on average a 5-fold increased IC50 dose compared to GBM CSCs, mouse astrocytic stem cells had a 7-fold increased IC50 dose, and human fibroblasts a 20-fold increased IC50 dose (**Fig. 6B, Table 1**). In order to directly compare the effect of C16 in non-malignant and malignant cells, we utilized a series of immortalized and/or transformed human neural stem cells from the same background (CB660 line) that are grown in identical media. These cells have been modified via the following combinations of loss of tumor suppressors and addition of oncogenes: dominant-negative p53^DD^ and hTERT (noted as “PhT”), CyclinD1 and CDK4^R24C^ (noted as “CC”), and MYC and H-RasV12, as previously described (Hubert et al., 2013). We found that sensitivity to C16 dramatically increased as a result of exogenous MYC expression, and most acutely (50- fold decrease in IC_50_) in MYC and H-RasV12 transformed cells, which are malignant (**Fig. 6C, Table 1**). These data demonstrate that sensitivity to WDR5 inhibition increases as a result of malignant transformation. It should be noted that the proliferation rate of immortalized neural stem cells increased with exogenous H-RasV12 and MYC expression, and we therefore wanted to determine whether increased sensitivity to the inhibitor was simply due to increased proliferation rate of cells. We determined doubling times for a panel of CSCs and transformed neural stem cells. We found no correlation between doubling times and C16 IC50s (**Supp. Fig. 6C**), suggesting the sensitivity to WDR5 inhibition is not solely due to proliferation rate. Based on these observations, we further tested the effects of C16 in a subset of CSC models (3832, DI318 and L0) and found that C16 reduced cell number over time and increased apoptosis in a concentration-dependent manner (**Fig. 6D,E, Supp. Fig. 6D,E**). C16 also reduced CSC proliferation in 4 independent patient-derived GBM organoids (**Fig. 6F**). Due to the observed growth inhibitory effect of C16 in human NSCs, we investigated further. Observation of these cells under the microscope indicated that these cells were undergoing a slowing of growth rather than cell death upon WDR5 inhibitor treatment, while cell death was evident upon WDR5 inhibitor treatment in GBM CSCs. To test this, we measured apoptosis in human NSCs in parallel with GBM CSCs treated with WDR5 inhibitor C16 for 4 days via annexinV staining by flow cytometry. We did not observe a significant increase in apoptosis in NSCs compared to GBM CSCs (**Fig. 6G**). Similarly, we did not observe a significant increase in apoptosis in primary astrocytes compared to GBM CSCs inhibition (**Supp. Fig. 6G**). These data demonstrate differential sensitivity of tumor vs. normal cells to WDR5 inhibition. Given these data, we interrogated WDR5 and WRAD complex expression via RNA sequencing data from a panel of normal brain cell types and GBM lines. These data show significantly increased expression of WDR5 and RBBP5 in GBM and a trend of increased expression of ASH2L and DPY30 (**Fig. 6H, Supp. Fig. 6F**). We compared protein expression of WDR5 and other WRAD complex members in GBM CSCs, NSCs and astrocytes and found diminished expression in normal brain cell types compared to GBM (**Fig. 6I)**. Likewise, WDR5 and RBBP5 protein expression also increases modestly in normal neural stem cells with the addition of Myc expression (**Supp. Fig. 6H**), which dramatically increased the cells’ sensitivity to pharmacologic WDR5 inhibition.

To assess *in vivo* toxicity of C16, we dosed mice with 10 mg/kg C16 daily and did not observe any reduction in weight or other phenotypic signs of potential toxicity over a 30-day period (**Supp. Fig. 7A**). At the end of this study, we collected organs and ran chemistry panels to further assess toxicity. We found no changes in liver enzymes, chloride, creatinine, sodium or glucose between control and C16-treated mice (**Supp. Fig. 7B**). Recently published work using C16-related WDR5 small molecule inhibitors also found no systemic toxicity and did not note any neurologic defects, noting a desirable oral pharmacokinetic profile with manageable intravenous clearance and high oral bioavailability (Teuscher et al., 2022). Mouse and human WDR5 are identical (Guarnaccia and Tansey, 2018), therefore C16 is predicted to bind mouse WDR5 and in this sense, toxicity studies with WDR5 inhibitors in mice are translatable to humans. However, assessment of brain penetration via a snapshot brain-to-plasma time course concentration profile after a single IP bolus dose of 10 mg/kg in CD-1 mice revealed limited brain penetration of C16, with an area under the curve brain-to-plasma ratio of less than 10% (AUC_brain_/AUC_plasma_ ratio <0.1) (**Supp. Fig. 7C**). C16 was submitted to Absorption Systems Inc. for *in vitro* determination of blood-brain-barrier penetration potential using MDR1-MDCK cell monolayers, a routine methodology used to predict the likelihood of passive CNS penetration. An average A-B passive permeability (Papp) of 0.715×10^-6^ cm/sec was measured for C16, a value that is indicative of low brain penetration potential classification. This *in vitro* observation is consistent with the observed *in vivo* AUC_brain_/AUC_plasma_ after a single IP administration. Typical CNS therapeutics maintain moderate to high passive permeability (P_app_ >10×10^-6^ cm/sec) and lack active transport/efflux from transporters expressed at the blood-brain barrier (Mahar Doan et al., 2002). Despite the expected low brain penetrance of C16, we tested C16 *in vivo* in the context of orthotopic brain tumors. NSG mice were intracranially implanted with DI318 CSCs and daily treatment of 10mg/kg C16 was started 10 days after implantation. As expected from the low drug penetrance through the blood-brain-barrier, we did not observe a difference in survival within the brain. Of note, the mice tolerated daily IP treatment of C16 well for almost 3 weeks in this experiment (**Supp. Fig 7D**).

Given these data, we implanted GBM CSCs as flank tumors rather than intracranially to test C16 *in vivo*. We found that both direct tumoral injection and systemic IP injection of C16 modestly reduced tumor volume (**Fig. 6J-K**). Mice were treated until vehicle group mice reached a humane endpoint based on flank tumor size. In these experiments, mice tolerated systemic injection of C16 for over 2 weeks without any side effects, supporting the idea that the inhibitor is tolerable and not overtly toxic to mice. Finally, in order to demonstrate that the tumor suppressive effect is related to WDR5, we prepared lysates from endpoint tumors (from **6K**) and performed co-immunoprecipitations to measure protein-protein interactions between WDR5 and its binding partner RBBP5. We saw a trend of reduction in WDR5 bound to RBBP5 (**Fig. 6L**), supporting a WDR5-specific effect in tumors after systemic IP injection of C16.

### WDR5 knockdown reduces CSC growth, self-renewal, and tumor initiation

To investigate the relevance of WDR5 in the context of human GBM patients, we interrogated publicly available gene expression data from patients’ tumors (Bowman et al., 2017). Importantly, WDR5 expression is elevated in GBM compared to normal brain (**Supp. Fig. 8A**). To examine WDR5 expression in more detail, we queried WDR5 and WRAD complex expression in RNA sequencing data through a recently reported “BRAIN-UMAP” (https://www.fredhutch.org/content/dam/www/research/divisions/human-biology/retreat-poster/poster-pdfs/Arora_Holland_Poster.pdf/ cite bioRxiv). The BRAIN-UMAP was generated by combining RNASeq abundance values from three different uniformly-processed pipelines for 702 adult glioma samples from The Cancer Genome Atlas (TCGA), 270 adult glioma samples from Chinese Glioma Genome Atlas (CGGA), 1409 healthy normal brain samples from Genotype-Tissue Expression Project (GTEx) across 13 GTEx-defined brain regions and 802 pediatric tumor samples from Children Brain Tumor Network (CBTN). UMAP projections of gene expression for WRAD complex members reveal enriched WDR5 and WRAD complex member expression in glioma samples from TCGA and CGGA compared to normal brain tissues from GTEx (**Fig. 7A,B; Supp. Fig. 8B, C**). In addition, increased WDR5 expression in patients’ tumors is associated with poorer overall survival in GBM (**Supp. Fig. 8D**) and correlates with increased SOX2 expression (**Supp. Fig. 8E**). Together, these data suggest WDR5 is important in patients’ tumors and may play a role in the SOX2+ CSC state.

**Figure 7:**
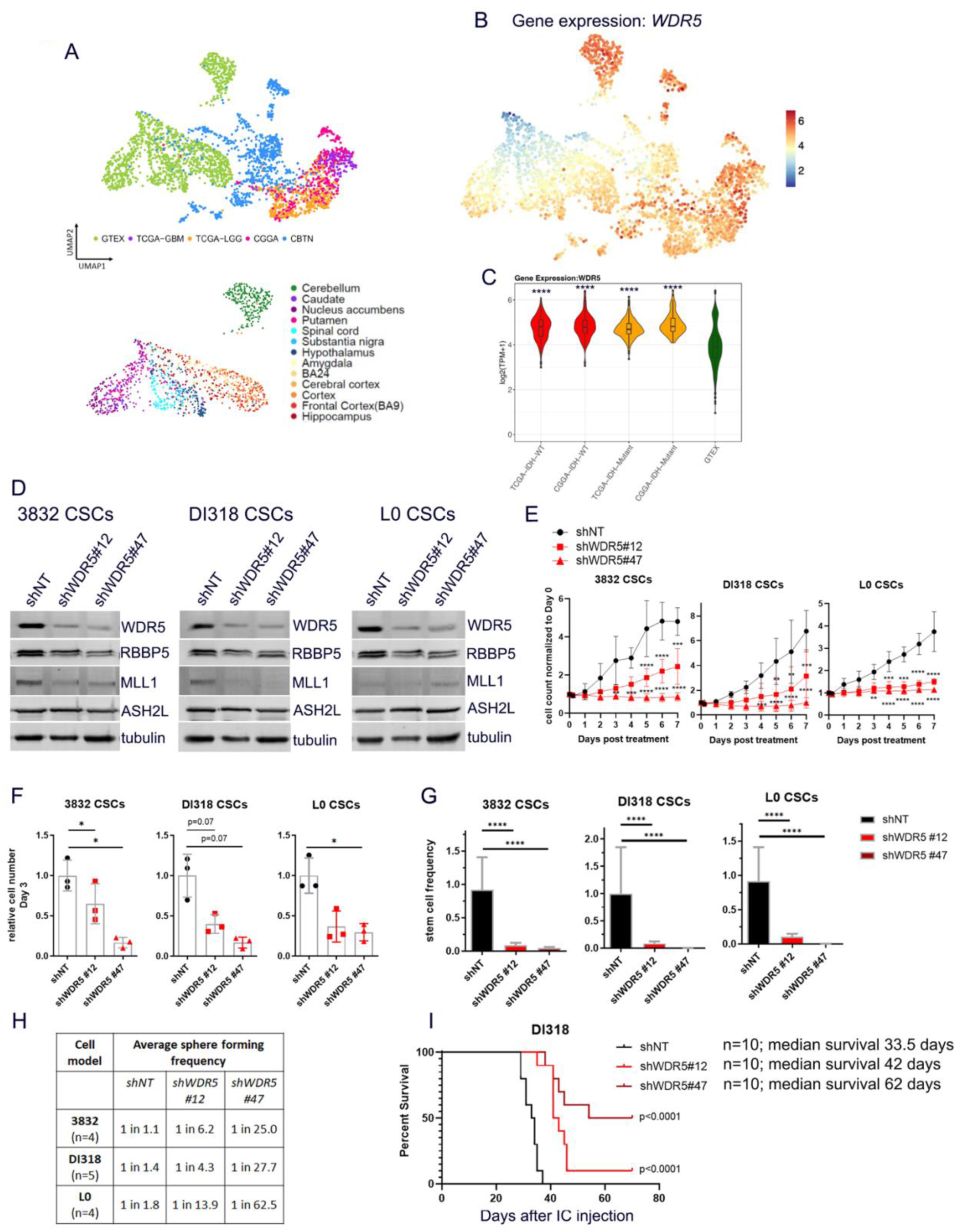
WDR5 knockdown reduces GBM CSC growth, self-renewal and tumor initiation. (**A**) UMAP projection showing brain tissue and tumor sample source. GTEX = brain tissue samples from Genotype-Tissue Expression Project; TCGA = The Cancer Genome Atlas; CGGA = Chinese Glioma Genome Atlas; CBTN = Children’s Brain Tumor Network. (**B**) UMAP projections of gene expression for WDR5. (**C**) RNA expression of WDR5 in selected tumor and normal groups. p values determined by unpaired t-tests with correction for multiple comparisons. (**D**) Short hairpin RNA-mediated targeting of WDR5 was done with 2 non-overlapping short hairpins in 3 CSC models, DI318, 3832 and L0. Western blots indicate the level of WDR5 protein in CSCs infected with a non-targeting (shNT) control virus or WDR5 knockdown (KD) viruses. (**E**) Proliferation of WDR5 KD and shNT control CSCs over 7 days, determined by Incucyte live cell imaging. Values represent mean fold change in cell count relative to Day 0, +/- standard deviation (SD), n=3 biological replicates, p values determined by two-tailed, unpaired t tests. (**F**) WDR5 KD and shNT control CSCs were plated, and viable cell counts were measured by CellTiter Glo luminescence viability assay after 72 hrs. Bars represent mean luminescence values relative to the average for shNT control cells, +/- SD, circles represent biological replicates, p values determined by one way ANOVA and post-hoc Dunnett’s multiple comparisons test. (**G**) *In vitro* limiting-dilution analysis was performed on WDR5 KD and shNT control CSCs. Bars represent sphere formation frequency from one representative experiment; error bars represent 95% confidence intervals for stem cell frequency. (**H**) Mean sphere formation frequency for each group is listed in the table. Independent biological replicates are shown in Supp. Fig. 8F. (**I**) Kaplan-Meier survival plot of mice intracranially implanted with WDR5 KD or shNT control CSCs. p values indicate comparisons between shNT and shWDR5 and were determined by log-rank analysis.

We finally sought to validate our inhibitor studies with WDR5 genetic loss of function. We silenced WDR5 using short hairpin RNA (shRNA) mediated interference (**Fig. 7D**) and found that WDR5 loss led to reduced expression of RBBP5 and MLL1, indicating that WDR5 inhibition may destabilize the WRAD complex. We next tested how WDR5 loss affected the stem cell behaviors of the 3832, DI318 and L0 CSC models. Depletion of WDR5 protein using shRNA resulted in reduced cell growth (**Fig. 7E, F**) and attenuated self-renewal (**Fig. 7G, H, Supp. Fig. 8F**) compared to non-targeting (NT) controls. As CSCs are functionally defined, in part, by their capacity for tumor initiation, we intracranially implanted CSCs expressing control or WDR5-targeted shRNAs and found that WDR5 knockdown increased tumor latency (**Fig. 7I**). In fact, half of mice implanted with shWDR5#47 CSCs, which leads to greater knockdown of WDR5, did not develop tumors within the timeframe of the study. These data imply that below a certain threshold of WDR5 expression, tumors are not viable. Taken together, these data more broadly validate our initial organoid screening results, demonstrate that WDR5 is essential for the CSC phenotype, and provide a rationale for the pharmacological targeting of WDR5 and the WRAD complex to impair CSC population viability in GBM.

## Discussion

The goal in our study was to determine if WDR5 is a viable therapeutic target in GBM. Our data provide proof of concept for developing WRAD inhibitors based on an unbiased screen, the use of a WDR5-directed tool compound, C16, and genetic loss-of-function studies on WDR5. Our findings demonstrate that the GBM CSC phenotype is reliant on WDR5 to a greater extent than non-stem tumor cells and non-malignant neural stem cell populations. We provide the first evidence for the role of WDR5 in regulating epigenetic maintenance of the GBM CSC state, which is consistent with our previous observation of the importance of one of its key binding partners, RBBP5, for GBM CSC self-renewal (Alvarado et al., 2017). Together with our observation of global reduction of H3K4me3 by western blot and CUT&Tag of C16 treated cells, we conclude that C16 causes a reduction in the H3K4me3 mark across the genome, particularly in key genes involved in neural developmental and oncogenic pathways, with a bias for POU domain motifs.

While H3K4me3 levels are documented to generally positively correlate with gene expression, after decades of research, it is still unclear how exactly H3K4me3 loss affects transcription. In our study, for some genes, we observed reduced expression when H3K4me3 was lost, but for other genes, such as the transcription factors EGFR, KLF8, SALL2, SOX4 and SOX5, there was minimal or no reduction in gene expression despite an observed reduction in H3K4me3. It is likely that the coordinated addition or removal of multiple chromatin modification changes is needed to significantly alter expression for certain genes. In addition, H3K4me3 loss was previously shown to result in minimal global transcriptional changes and H3K4me3 has been shown to be dispensable for maintenance of transcription (Howe et al., 2017; Murray et al., 2019). Evidence from multiple studies suggest different roles for this mark, including regulation of alternative splicing, regulation of miRNA genes, transcriptional memory and stabilization of transcriptional noise (Guenther *et al*., 2005; Howe *et al*., 2017). Regardless of its specific role, our studies show that WDR5 inhibitor-mediated H3K4me3 loss compromises CSC survival. In future studies, we aim to determine the transcriptional impact of H3K4me3 loss in GBM CSCs, perhaps through matched single cell H3K4me3 and transcriptional profiling.

Moreover, our findings align with previous observations of the importance of MLL1, which is dependent on WDR5 for its function, in GBM CSCs (Gallo et al., 2013; Heddleston et al., 2012),. Notably, MI-2-2, a menin–MLL inhibitor, inhibited temozolomide-resistant GBM clones and reduced subcutaneous GBM growth *in vivo* (Lan et al., 2017). WDR5 loss led to global reduction of H3K4me3 in various cell models (Benayoun *et al*., 2014; Siladi *et al*., 2022), as did the WDR5 inhibitor Piribedil (Zhang et al., 2018), yet other studies with WDR5 inhibitors did not always observe this phenomenon. As mentioned, C16 targets the WIN site of WDR5, and this WIN site interaction is only necessary for the activity of MLL1 and SET1A and not other SET1 family methyltransferases. In addition, optimal histone methyltransferase activity of the MLL1 complex depends on WDR5, while the other SET1-family methyltransferases can be fully activated by just RBBP5 and ASH2L (Alicea-Velazquez *et al*., 2016; Li *et al*., 2016; Patel *et al*., 2008). Therefore, in theory, WDR5 WIN site inhibition should primarily affect MLL1 function. However, our observation of global diminishment of H3K4me3 after C16 treatment may suggest that C16 has an effect broader than MLL1, since MLL1 has been previously demonstrated to have proportionally fewer targets than other methyltransferases in the family. Specifically, MLL1 has been documented to only deposit H3K4me3 at specific genomic loci, in contrast to SET1A/SET1B which are credited with genome-wide deposition of this mark (Sze et al., 2020). Additionally, in previous studies, MLL1 knockout mouse embryonic fibroblasts only displayed H3K4me changes in ∼5% of promoters analyzed by ChIP-chip (Wang et al., 2009). However, as the authors state, this could be attributed to compensation of methyltransferase activity by other SET1 family members. Since the loci targeted by MLL1 in GBM have not been previously examined, a possibility is that MLL1 methylates a broader and/or different array of targets in GBM than in the cell types previously examined, primarily leukemia, embryonic stem cells and fibroblasts. For instance, MLL1 is well known to regulate *HOX* genes, yet we did not observe downregulation of H3K4me3 on *HOX* genes in our GBM models. Alternatively, given C16 treatment (WIN site inhibition) led to disruption of the non-WIN site mediated interaction between WDR5 and RBBP5, it is likely that WDR5 functions beyond those mediated through MLL1 are affected by this inhibitor. This may explain why our system displays global H3K4me3 loss, whereas in other systems (other WDR5 WIN site inhibitors and MLL1 knockdown), H3K4me3 levels are not globally altered. Investigation into specific roles of WDR5 in various tumor models is thus an important avenue for future study.

Identifying mechanisms to attenuate the CSC state remains an immediate priority for malignant cancers, including GBM, and it is well established that self-renewal is driven by the coordinate action of transcription factors and programs (Mehta et al., 2011; Rheinbay et al., 2013; Suva *et al*., 2014). There have been recent promising efforts to target individual transcription factors, such as with the OLIG2 inhibitor CT-129 (Oasa et al., 2020), but DNA/protein interactions have been historically difficult to target (Bushweller, 2019). Therefore, understanding the upstream molecular network of self-renewal transcriptional programs may provide more rational therapeutic targets. Tumor cells in a variety of cancers have shown dependence on WDR5 for survival, including in leukemia, pancreatic cancer, breast cancer, and ovarian cancer (reviewed in (Aho et al., 2019b; Lu et al., 2018; Zhao et al., 2021). The role of WDR5 appears to vary in different cancers, likely due to the array of WDR5 interaction partners that have been identified (Guarnaccia *et al*., 2021). Likewise, WIN site inhibitors appear to have different mechanisms of action across tumor types. Perhaps the best studied role of WDR5 is in MLL-rearranged AML, where one WIN site inhibitor led to global H3K4me2/3 reduction (Zhang et al., 2018), while another led to H3K4me3 reduction specifically on HOX genes (Cao *et al*., 2014). Yet another report found that WDR5 WIN site inhibitors led to potent induction of apoptosis in MLL1-rearranged AML by obstructing protein synthesis capacity, independently of changes in histone methylation (Aho *et al*., 2019a). Here, we link WDR5 WIN site inhibition in GBM to reduced H3K4me3 on genes involved in pathways previously shown to be important for driving CSCs in GBM, including the WNT and EGFR pathways (Furnari *et al*., 2015; Kim et al., 2013). Together, these data suggest a context- and tumor-type specific role of WDR5, likely underlain by differences in expression and function of binding partners, available chromatin binding sites, and downstream signaling networks. In our study, the inhibition of WDR5 appears to be driving a global reduction in the H3K4 methyltransferase activity of the WRAD complex, which represents a vulnerability to tumor populations as compared with non-malignant control cells, however, it still remains unclear exactly how C16, the WDR5 WIN site inhibitor tested in the current study, leads to cell death. This represents an immediate future direction as it may also provide insight into putative therapeutic resistance mechanisms. Our screen identified WDR5 as essential in the SOX2+ organoid niche, but other WRAD complex components did not come out of the screen, indicating that WDR5 likely has additional, WRAD-independent roles in GBM CSCs. Due to the nature of screens where efficiency of knockdown from gene to gene can vary, it is also possible that other WRAD complex members and MLL1 may have been included as false negative hits in our screen. However, we propose that the most likely explanation for our results is that WDR5’s role in the WRAD complex combined with its other functions results in a stronger phenotype upon WDR5 knockdown than knockdown of individual components of the WRAD complex. Given the versatile nature of WDR5 and its binding partners and downstream targets across cancer types, a more focused assessment in each cancer, including GBM, is warranted. In addition, the variation in response to WDR5 inhibitors in GBM CSC models demonstrates that some patient tumors may be more sensitive and identifies a need to find biomarkers associated with increased sensitivity.

A current limitation is the limited brain penetration of C16, which reveals a clear need for medicinal chemistry efforts to surmount challenges within the current existing scaffolds (such as poor permeability and potential transporter efflux) and turns attention to inhibitor modifications that will lead to overall improved brain penetration. Our transformed neural stem cell models and elevated expression of WDR5 in GBM compared to normal brain suggest the existence of a therapeutic window to target WDR5 in CSCs without compromising normal neural function, and future drug developments must take care to maintain this window. Data from DepMap portal (Broad Institute) (Dempster et al., 2019; Ghandi et al., 2019; Meyers et al., 2017) demonstrate that WDR5 is “common essential” in large, pan-cancer screens, indicating that WDR5 inhibitors could be broadly applicable in multiple tumor types. Taken together, our findings provide the first report linking WDR5 to the GBM CSC phenotype, highlight a key role for WDR5 in the epigenetic maintenance of the CSC state, and provide a starting point for WDR5 inhibitors to neutralize CSC populations in GBM and potentially other advanced cancers.

## Acknowledgements

We thank members of the Lathia, Hubert, Stauffer, Paddison and Patel laboratories for critical feedback. We also thank Ms. Amanda Mendelsohn for illustration assistance. This work was supported by National Institutes of Health (NIH) T32CA059366 and NIH F32NS116109 to K.M.; NIH F30CA250254 to A.J.L.; NIH/NCATS CTSA KL2TR0002547, the Clinical Translational Science Collaborative of Cleveland grant UL1TR002548, an American Brain Tumor Association Discovery Grant in memory of Dr. Joseph Weiss, grant #IRG-16-186-21 to the Case Comprehensive Cancer Center from the American Cancer Society, and support from the VeloSano Cancer Research Fund to C.G.H.; R35 CA197718 to J.N.R.; and the Lerner Research Institute and Case Comprehensive Cancer Center to J.D.L. Work in the Lathia laboratory is also supported by NIH grants P01 CA245705 and R35 NS127083. S.R.S thanks the Lerner Research Institute for lab startup support and Cleveland Clinic Philanthropy and an anonymous donor for a generous donation to support capital purchases for this research. We thank the Vanderbilt University Medical Center Metabolic Physiology Shared Resource Core and the Diabetes Research and Training Center for in-life brain plasma studies using compound C16. We thank Alan Pratt and the Prevention Research Laboratory and Lab Diagnostic Core at Cleveland Clinic for blood chemistry studies using compound C16. The Metabolic Physiology Shared Resource is supported by NIH grant DK020593.

## Author Contributions

K.M., C.G.H., J.D.L., S.R.S., J.N.R. & J.A. designed the study and experiments; K.M., S.S., S.A.S., D.J.S., C.M.G., L.W., G.R., R.S., K.K., C.G.H., A.L., J.D.K., A.K., S.J. & J.A. performed experiments and analyzed data; T.E.M. provided guidance on the organoid screen; A.P.P & P.J.P. provided guidance on CUT&Tag experiments and data analysis. S.A., K.M. & J.B. performed statistical analyses. K.M., C.G.H., J.D.L., S.R.S. & E.E.M-H. wrote the manuscript. All authors approved the final version of the manuscript.

## Methods

### RESOURCE AVAILABILITY

#### Lead contact

Further information and requests for resources and reagents should be directed to and will be fulfilled by the lead contact, Justin D. Lathia.

#### Materials availability

This study did not generate new unique reagents.

### EXPERIMENTAL MODEL AND SUBJECT DETAILS

#### Animals

For flank and intracranial tumor experiments, NSG mice (NOD.Cg-*Prkdc^scid^ Il2rg^tm1Wjl^*/SzJ; stock 005557; Jackson Laboratory) were bred in house (Cleveland Clinic). For DI318 intracranial tumor experiments, 7.5-week-old NSG mice were used. For DI318 flank tumor experiments, NSG mice at least 8 weeks of age were used. For L0 flank tumor experiments, 8-week-old NSG mice were used. An equal number of male and female mice were used for all animal experiments and were evenly distributed between experimental groups. For the IP dosing weight study, 8-week-old NSG mice were used. Mice were housed in the Cleveland Clinic Biological Resources Unit. Mice maintained on a 12-hour light cycle (0600-1800). Room temperature was monitored daily and maintained at 22-25°C. All experiments were performed in compliance with institutional guidelines and were approved by the Institutional Animal Care and Use Committee of the Cleveland Clinic (protocol 2019-2195).

#### Primary cell cultures

Cancer stem cell (CSC) models were generated by passaging primary tumor cells as GBM xenografts as previous described (Lathia et al., 2010). Briefly, primary tumor cells were intracranially implanted into NSG mice, and upon tumor formation, tumors were isolated, digested with papain (Worthington) as described previously (Alvarado et al., 2017), and dissociated cells were plated overnight in Neurobasal™ medium minus phenol red (Gibco) with 1X B-27 supplement (Gibco), 1 mmol/L sodium pyruvate, 2 mmol/L L-glutamine, 50 U/mL penicillin/streptomycin, 20 ng/ml human (h)EGF and 20 ng/ml hFGF2 (R&D systems). Subsequently, CD133+ cells were isolated by magnetic bead sorting (Miltenyi). CD133+ cells were cultured in the media described above. Some cell models were previously established at other institutions (**Table 1**). CD133+ cells were seeded in suspension culture at 5×10^4^ cells/ml and passaged no more than 10 times. After 10 passages, cells were re-implanted into NSG mice and enriched for CD133+ cells.

Organoids were formed as previously described (Hubert *et al*., 2016) by suspending tumor cells in 80% Matrigel (BD Biosciences) and forming 20 μl pearls on parafilm molds prior to culture. Organoids were seeded with 10,000 cells per organoid and cultured in 6-well or 10-cm plates with shaking in supplemented Neurobasal media as above with the addition of phenol red. Transformed human neural stem cells (CB660) were generated as previously described (Hubert *et al*., 2013). These include NSC-CB660 cells with dominant-negative p53DD and hTERT (PhT); NSC-CB660 cells with dominant-negative p53^DD^, hTERT, CyclinD1 and CDK4^R24C^ (PhTCC); NSC-CB660 + PhTCC + Myc; NSC-CB660 + PhTCC + H-RasV12; and NSC-CB660 + PhTCC + Myc + H-RasV12. These lines were grown adherently on plates coated with 10 µg/ml laminin (Sigma) and cultured in a 1:1 ratio of DMEM-F12 and NeuroCult NS-A Basal Medium (Human) (Stem Cell Technologies) with 1X N-2 supplement (Gibco), 1X B-27 supplement, sodium pyruvate, L-glutamine, 1X pen/strep, 20 ng/ml human (h)EGF and 20 ng/ml hFGF2. Cells were grown at 37°C with 5% CO2. The sex of the cells is as follows: 3832 (female), DI318 (male). The sex of other GBM models is not known. De-identified GBM specimens were collected from the Cleveland Clinic Brain Tumor and Neuro-Oncology Center in accordance with an Institutional Review Board-approved protocol, and informed consent was obtained from all GBM patients contributing tumor specimens.

### METHOD DETAILS

#### Organoid screen

To investigate the effects of targeting epigenetic regulators in glioblastoma cells within the tumor microenvironment, we used an inducible RNAi screening system that was previously used for screening *in vivo* (Miller et al., 2017). Our shRNA library contained 1,586 shRNAs targeting 406 known chromatin and transcriptional regulator genes (2–4 shRNAs per gene), with positive and negative control shRNAs. GBM528 patient-derived CSCs were transduced with the shRNA library pool at low MOI to ensure single viral integration, and cells with genomic integration of shRNAs as monitored by expression of a constitutive mVenus fluorescent reporter were isolated using FACS. Cells were allowed to recover and expand for three passages.

Organoids for shRNA screening were formed as described above by seeding 30,000 positively infected cells per 20 µl organoid using custom 96-well-format parafilm molds and multichannel pipettes. Organoids were allowed to grow and mature uninduced for 1 month in 500 ml spinner flasks (Corning #3578) in 250 mL of media at 37°C with 5% CO2, prior to the addition of doxycycline (1 µg/ml, Sigma-Aldrich) to induce gene knockdown. Organoids were maintained on doxycycline for 21 days. Each of 3 organoid screen cohorts, consisting of 55 independent organoids each, were processed and analyzed separately. This represents an approximate 1000-fold library coverage for each screen replicate. Day 0 controls were also collected, stored frozen, and processed in parallel for comparison. At the end of the screen, organoids were regionally labeled with CellTracker CMAC (Molecular Probes) for 2 hrs as previously described *(Shakya et al.*, 2021), and single organoids from each screen cohort were spot-checked by confocal microscopy in a compatible dish (MatTek 35 mm glass bottom dish P35G-1.5-10-C) to ensure proper CMAC labeling. Labeled organoids were dissociated and separated by FACS sorting, marked by positivity for constitutive mVenus expression and doxycycline-induced dsRed expression, and sorted into regional populations based upon retention of CMAC regional blue dye.

Genomic DNA was isolated from each screened population and sequenced as described (Miller et al., 2017). Genomic DNA was isolated by two rounds of phenol extraction using PhaseLock tubes (5prime) followed by isopropanol precipitation. Deep sequencing libraries were generated by PCR amplification of shRNA guide strands using barcoded primers that tag the product with standard Illumina adapters (p7+loop, 5′-CAAGCAGAAGACGGCATACGA-NNNN (4 nucleotide barcode)-TAGTGAAGCCACAGATGTA-3′; p5+miR3′, 5′-AATGATACGGCGACCACCGATGGATGTGGAATGTGTGCGAGG-3′). Libraries were sequenced on the HiSeq 2500 platform at the Cleveland Clinic Genomics Core Facility. Libraries were sequenced using a primer that reads in reverse into the guide strand (miR30EcoRISeq, 5′-TAGCCCCTTGAATTCCGAGGCAGTAGGCA-3′). Sequence processing was performed as previously described using two custom workflows at https://usegalaxy.org. Raw read counts were converted to reads per million (RPM) to control for variations in total shRNA reads in each sample. shRNAs were scored using RIGER and extension of the GENE-E package (Broad Institute) (Luo et al., 2008). Median RPM value for each replicate was used for analysis. The signal-to-noise ratio of replicates was used to calculate individual shRNA score based on their ability to deplete cells in the induced cohorts compared to the control inputs, and second - best shRNA score was used to rank genes. Expressed genes with a total RIGER p-value score <0.05 for depletion compared to controls were considered hits.

#### Organoid IHC for SOX2/pHH3

Organoids were treated with drugs as indicated while shaking in 6-well plates. Treated organoids were then fixed in 10% neutral buffered formalin for at least 24 hrs prior to transfer to 70% ethanol and subsequent paraffin embedding by the LRI Biomedical Engineering histology core. Sections (4 µm) were cut, placed on slides, deparaffinized, unmasked by boiling in 1X citrate solution (Cell Signaling) and blocked with normal donkey serum or BSA. Antigens were detected using anti-SOX2 (R&D, #AF2018) and anti-phospho-Histone H3 (Cell Signaling, #9701S) antibodies. Detection was performed with DAB and counterstained with Gills 2 Hematoxylin and bluing reagent. Coverslips were mounted with Permount, and whole slides were scanned on a Leica Aperio AT2 digital slide scanner using a 20X objective in the LRI imaging core. For immunofluorescence, DAPI (1:10,000) was used for DNA detection and images were acquired with the Leica DM5500B upright microscope and Leica DFC 7000 GT monochrome camera (Leica Biosystems). Image fields were extracted using Leica ImageScope software.

#### Chemical synthesis & WDR5 time-resolved fluorescence energy transfer (TR-FRET) competition assay

Compound 16 (C16) was synthesized as previously published (Tian et al., 2020). Recombinant His6-SUMO-WDR5 was expressed and purified as previously published (Tian et al., 2020). The WDR5 TR-FRET Competition Assay was run following previously published methods (Tian et al., 2020). C16 was tested for MLL1-FITC probe displacement using a 10-point CRC with a top concentration of 10 µM and 5-fold dilution scheme. The 520/495 FRET ratio was plotted against compound concentration and fit with a “One Site – Fit Ki” in PRISM 8, with “HotNM” constrained to 150 nM and the “HotKdNM” constrained to 2 nM. C16 was tested in three independent experiments with duplicates run for each experiment (n=6 total).

#### Western blotting & co-immunoprecipitation

For protein isolation, cells were washed out of medium with PBS. Lysates were prepared using modified radioimmunoprecipitation assay (RIPA) buffer containing protease and phosphatase inhibitors (50 mM Tris-HCl, pH 7.4, 1% NP-40 (vol/vol), 0.25% Na-deoxycholate (wt/vol), 150 mM NaCl, 1 mM EDTA, 1X Sigma p8340 Protease Inhibitor Cocktail, Sigma p5726 Phosphatase Inhibitor Cocktail, 1 mM NaF, 1 mM PMSF). Cells were lysed for 30 min on ice and centrifuged at maximum speed in a tabletop centrifuge to remove debris. Protein concentration was measured on a spectrophotometer (read at 595 nm) using Bradford reagent (500-0006; Bio-Rad). SDS-PAGE was performed, and cell lysates were resolved on polyacrylamide gels. Proteins were transferred onto PVDF membranes and blocked with TBST+5% BSA. A ChemiDoc MP imaging system (Bio-Rad) was used for visualization. For co-immunoprecipitation experiments, lysates were prepared as described above. Protein lysate (500 µg) was incubated with 5 µg immunoprecipitation antibody at 4°C overnight with rotation followed by incubated with protein A/G agarose beads for 1 hour at 4°C with rotation. Beads were washed 5 times with RIPA buffer, and bead-bound proteins were isolated by boiling antibody-bead complexes in SDS sample buffer. Immunoblotting was performed as described above. For C16 treatment western blots and co-immunoprecipitation, cells were plated at 5×10^5^ cells/ml with the indicated concentrations of inhibitor.

#### CUT&Tag

DI318 CSCs were treated for 72 hrs with 3μM C16. Cells were harvested, counted and CUT&Tag-IT Assay and library preparation was performed on 500,000 cells per replicate, according to the Manufacturer’s Manual (Active Motif). Rabbit anti-human Tri-Methyl-Histone H3 (Lys4) (C42D8) (Cell Signaling Technologies 9751) antibody was used for the CUT&Tag procedure. Size distribution and concentration of libraries was assessed using an Agilent 4200 TapeStation with D1000 reagents and Qubit Assay. Barcoded libraries were mixed to achieve equal representation and paired-end Illumina sequencing was performed on the barcoded libraries on a NovaSeq SP100 with the following parameters Read1:i7:i5:Read2=28:10:10:90.

#### Real-Time Reverse Transcription Polymerase Chain Reaction

RNA was isolated from cells using the Direct-zol RNA miniprep kit (Zymo Research). cDNA was synthesized with Superscript IV First Strand Synthesis System using dT primers (Invitrogen). qPCR was performed using SYBR-Green Mastermix (SA Biosciences) on a Viia7 () system using primers listed in the resources table below. Ct values for each gene were normalized to Actin levels and to DMSO treated cells.

#### SORE6-GFP reporter experiments

For SORE6-GFP reporter experiments (SOX2/OCT4 promoter response elements tagged to destabilized GFP), SORE6-dsCopGFP lentiviral particles were generated by transfection of 293T cells. 293T cells were transfected (Fugene transfection reagent) with pPACKH1 vectors and SORE6-dsCopGFP plasmid DNA (kindly provided by Wakefield Lab, NIH) according to the manufacturer’s protocols (System Biosciences). Viral supernatant was collected at 48 hrs, and virus was concentrated with PEG-it Virus Precipitation solution (System Biosciences). SORE6-GFP virus was added to CSCs plated on Geltrex. Forty-eight hrs after infection, 2-3 µg/ml puromycin was added to cells. After puromycin selection, cells were collected, and GFP^high^ (10-20% brightest) or GFP^negative^ cells were isolated by FACS. GFP^high^ and GFP^negative^ cells were subjected to limiting dilution analysis as described above, or protein was isolated for western blot. GFP^high^ cells were cultured further and used for inhibitor treatment experiments. Fluorescence images were taken with the IncuCyte Live Cell Analysis System (Sartorius).

#### Limiting dilution analysis

Cells were plated at 100 cells per well in 12 wells of a 96-well plate, and two-fold serial dilutions were performed. Twelve wells of each cell dose were plated. Limiting-dilution plots and stem-cell frequencies were calculated using ELDA analysis (http://bioinf.wehi.edu.au/software/elda/index.html; (Hu and Smyth, 2009)). For LDAs with C16 treatment, cells were incubated with inhibitor for the duration of the experiment.

#### IC_50_, cell growth, viability, apoptosis

Inhibitors were reconstituted to 10 mM in DMSO. For IC_50_ determination, cells were plated at 20,000 cells/ml in Geltrex-coated 96-well plates (to promote adherence) and treated with a 9 point, 2- or 3-fold serial dilution of inhibitor. For IC_50_ calculations, normalization was performed relative to the DMSO condition (100%) and a well with no cells (0%). After 7 days, cell viability was determined by ATP quantification with the CellTiter-Glo® Luminescent Cell Viability Assay (Promega). For cell growth assays, cells were plated at 20,000 cells/ml in Geltrex-coated 96-well plates and treated with different doses of inhibitor, then imaged using the IncuCyte Live Cell Analysis System using the cell-by-cell module (Sartorius). For apoptosis assays, cells were plated in duplicate at 20,000 cells/ml in Geltrex-coated 96-well plates and treated with different doses of inhibitor. Caspase 3/7 activity was determined with the Caspase-Glo 3/7 assay (Promega), and caspase activity was normalized to cell number by performing the CellTiter Glo Luminescent Cell Viability Assay on the duplicate plate. For quantification of apoptosis over time, cells were plated at 20,000 cells/ml in Geltrex-coated 96-well plates and treated with different doses of inhibitor in the presence of 1:1000 IncuCyte® Caspase-3/7 Dye for Apoptosis (Sartorius). Doubling times were calculated by determining cell counts over multiple days with the IncuCyte Live Cell Analysis System cell-by-cell module (Sartorius).

#### BBB penetration potential using MDR1-MDCK cell monolayers

MDR1-MDCK cell monolayers were grown to confluence on collagen-coated microporous membranes in 12-well assay plates. The permeability assay buffer was Hanks’ balanced salt solution containing 10 mM HEPES and 15 mM glucose at a pH of 7.4. The buffer in the receiver chamber also contained 1% bovine serum albumin. The dosing solution concentration was 5 μM of test article in the assay buffer. Cell monolayers were dosed on the apical side (A-to-B) or basolateral side (B-to-A) and incubated at 37°C with 5% CO_2_ in a humidified incubator. Samples were taken from the donor and receiver chambers at 120 minutes. Each determination was performed in duplicate. All samples were assayed by LC-MS/MS using electrospray ionization. Further details can be found at Absorption.com, assay #EA203.

#### *In vivo* brain:plasma study in mice

C16 was formulated from powder as 2 mg/mL solution in a 20% 2-(hydroxypropyl)-β-cyclodextrin in ddH_2_O (HP-β-CD; Sigma, catalog #C0926) solution. The solution was then made acidic with 1.0 equivalent of aqueous 1N HCl. The mixture was vortexed briefly and then sonicated for 5 min in a room temperature water bath sonicator to afford a clear solution to fine microsuspension. Animals were injected with a maximal dosing volume of 5 mL/kg to give a final 10 mg/kg body weight dose.

Male CD-1 mice (Charles River Laboratories, Wilmington, MA) were overnight fasted on the evening prior to study (food removed between 1500-1600 h). On the morning of study mice were weighed and allowed to acclimate to the room for at least 30 min prior to dosing. Food was returned 3 hrs after injection. At time 0, an IP injection of C16 was given. At 0.5 h, 1 h, 3 hrs, and 6 hrs after injection (n=2 per time point), mice were placed into a plane of anesthesia using Isoflurane. A terminal blood sample was collected via cardiac puncture followed by immediate euthanasia and brain collection. Brain was washed with cold PBS or Saline, blotted dry on a piece of gauze, weighed, and flash frozen in liquid Nitrogen. Whole blood was centrifuged at 5000-6000 g for 5 minutes and plasma was removed into a fresh tube for storage. All samples were stored at −80°C until shipment on dry ice to Q2 Solutions for tissue distribution bioanalysis (Q2 Solutions Bioanalytical and ADME Laboratories, Indianapolis, IN). The plotted time-course exposure plot for C16 represents the average concentrations of processed brain and plasma samples (brain homogenate supernatant and plasma) as determined by LC-MS/MS.

#### WDR5 knockdown

MISSION® pLKO.1-puro Non-Mammalian shRNA (SHC002) and WDR5 knockdown plasmids were purchased from Sigma. Several clones were tested, and 2 non-overlapping clones with efficient knockdown were selected to produce lentiviral particles (TRC clone IDs: TRCN0000157812 (shWDR5#12) and TRCN0000118047 (shWDR5#47)). For virus production, pLKO.1-shRNA plasmids were transfected into 293T cells along with psPAX and pMD2.G packaging plasmids to produce lentivirus. Forty-eight and 72 hrs after transfection, supernatant containing lentiviral particles was collected and concentrated with PEGit virus precipitation solution according to manufacturer’s protocol (System Biosciences). CSCs were plated on Geltrex, and virus was added to culture medium (MOI = 2), and then selected with 2-4 µg/ml puromycin.

#### Intracranial implantation

Intracranial tumor implantations were performed as described previously (Bayik *et al*., 2020). NSG mice were anesthetized with inhaled isoflurane for the duration of the procedure. For shRNA experiments, a total of 10,000 DI318 CSCs infected with control or WDR5 shRNAs were suspended in 10 µl Neurobasal null medium and stereotactically implanted in the left hemisphere ∼2.5 mm deep into the brain. For drug treatment experiments, a total of 5,000 DI318 CSCs were implanted intracranially into mice, and 10 days later, 10 mg/kg C16 was injected IP daily (formulated as described in “*In vivo* brain:plasma study in mice” section. Mice were monitored for neurologic signs and weight loss and deemed at endpoint when exhibiting any of these symptoms.

#### Flank tumor experiments

NSG mice were implanted subcutaneously with 500,000 DI318 or L0 human GBM CSCs. After tumor formation (3 weeks for DI318, 10 weeks for L0), 3 mg/kg C16 was injected daily directly into the tumors or 10 mg/kg C16 was injected daily intraperitoneally. For intratumoral dosing, C16 was dissolved at 5.1 mg/ml in 17.5% DMSO in PBS and treatment was started 3 weeks after tumor cell injection when tumors reached a volume of ∼100mm^3^. Tumor volume was calculated using the following formula for ellipsoid volume: 4/3π(w/2)^2^(h/2). For intraperitoneal (IP) dosing, C16 was dissolved at 2 mg/ml in 20% hydroxypropyl beta cyclodextran (BCD) in ddH2O. The solution was then made acidic with 1.0 equivalent of aqueous 1N HCl. Treatment was started 10 weeks after tumor cell injection when tumors reached a volume of ∼500mm^3^. When any animals in the experiment reached endpoint (determined by tumor size), mice were euthanized.

### QUANTIFICATION AND STATISTICAL ANALYSIS

Western blot quantification was performed using ImageJ (v1.53k, National Institutes of Health). For two group comparisons, P values were calculated using unpaired or paired two-tailed t tests. For multiple group comparisons, one-way ANOVA with post hoc tests were used as indicated in the figure legends. Log-rank tests were used for survival analysis. GraphPad Prism 9 was used for statistical tests. All *in vitro* experiments were done in technical triplicates for each experimental group, and multiple independent experiments were performed. To determine the number of mice needed per group for animal experiments, we utilized the Guidelines for the Care and Use of Mammals in Neuroscience and Behavioral Research from the National Research Council to estimate the minimal number necessary to achieve statistical significance (p < 0.05) for all tumor growth studies. The number of animals per arm was based upon the following calculation: = (1 + 2*C*)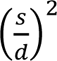, where *n* = Number of Animals per Experimental Group; *C* = 9.18 when α = 0.05 and 1 – β = 0.85 (Significance level of 5% with a power of 85%); *s* = Standard Deviation (≈ 7 days); *d* = Difference to be Detected (≥ 10 days). Thus, *n* = 10 animals were used per group, and to control for sexual dimorphism, males and females were treated as separate experimental groups and combined if there were no differences in the measured outcomes. *n* represents independent experiments (biological replicates) or individual mice. Statistical details can be found in figure legends. p<0.05 was considered statistically significant. *, p < 0.05; **, p < 0.01; ***, p < 0.001; ****, p < 0.0001.

#### CUT&Tag bioinformatic analysis

CUT&Tag reads were aligned to the human genome (hg38) using Bowtie2 (Langmead and Salzberg, 2012) as previously described (Henikoff *et al*., 2020). MACS2 was used for peak calling (Zhang et al., 2008) and peaks were annotated using ChIPseeker (Yu et al., 2015). BedTools (Quinlan and Hall, 2010), DiffBind (Ross-Innes *et al*., 2012) and DESeq2 (Love *et al*., 2014) were utilized to identify unique peaks, consensus peaks and perform differential analysis between groups (with significance set to False Discovery Rate (FDR) < 0.05). Fastq files and narrow peak files for each sample were deposited in GEO (GSE199110).

### RESOURCES TABLE

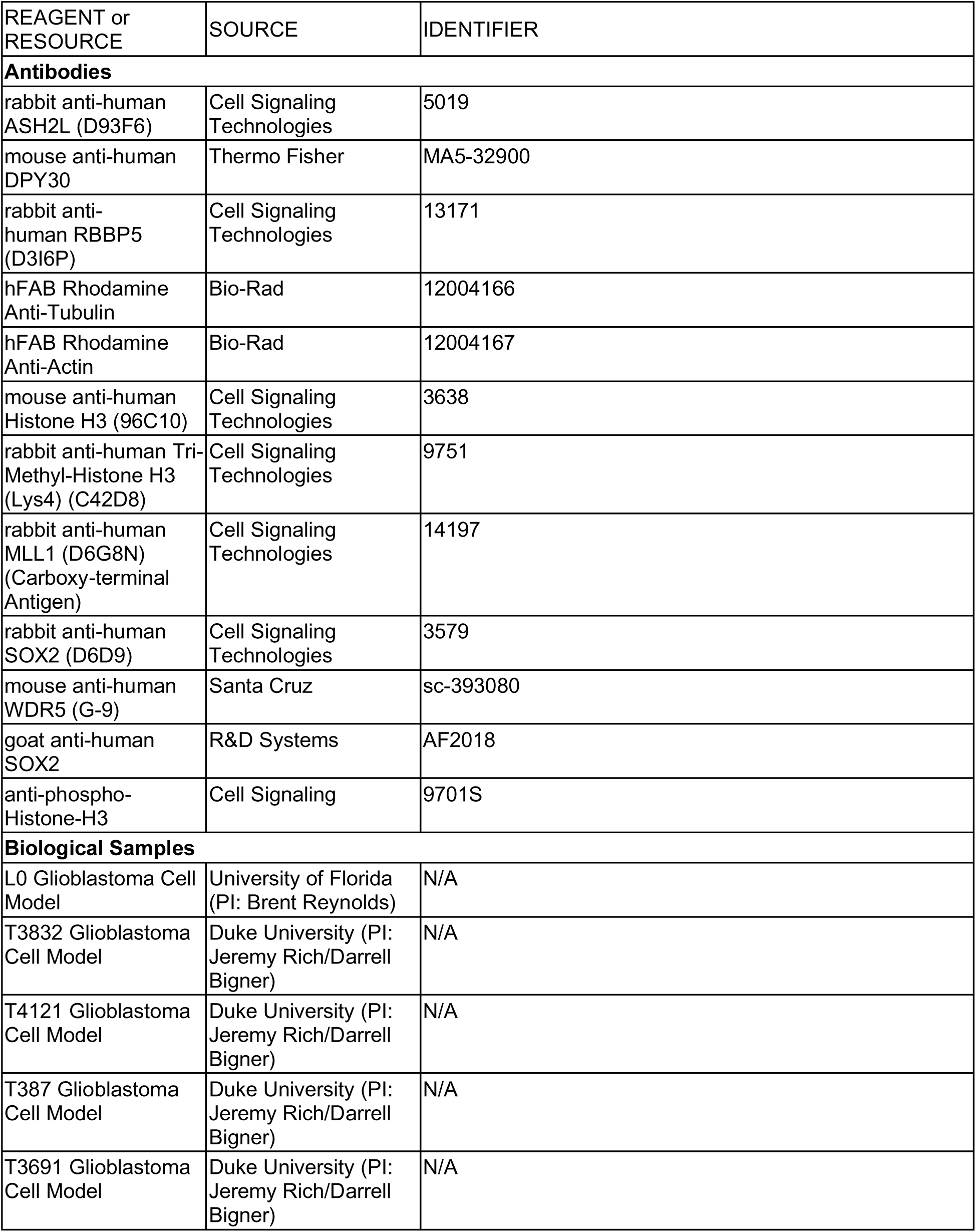
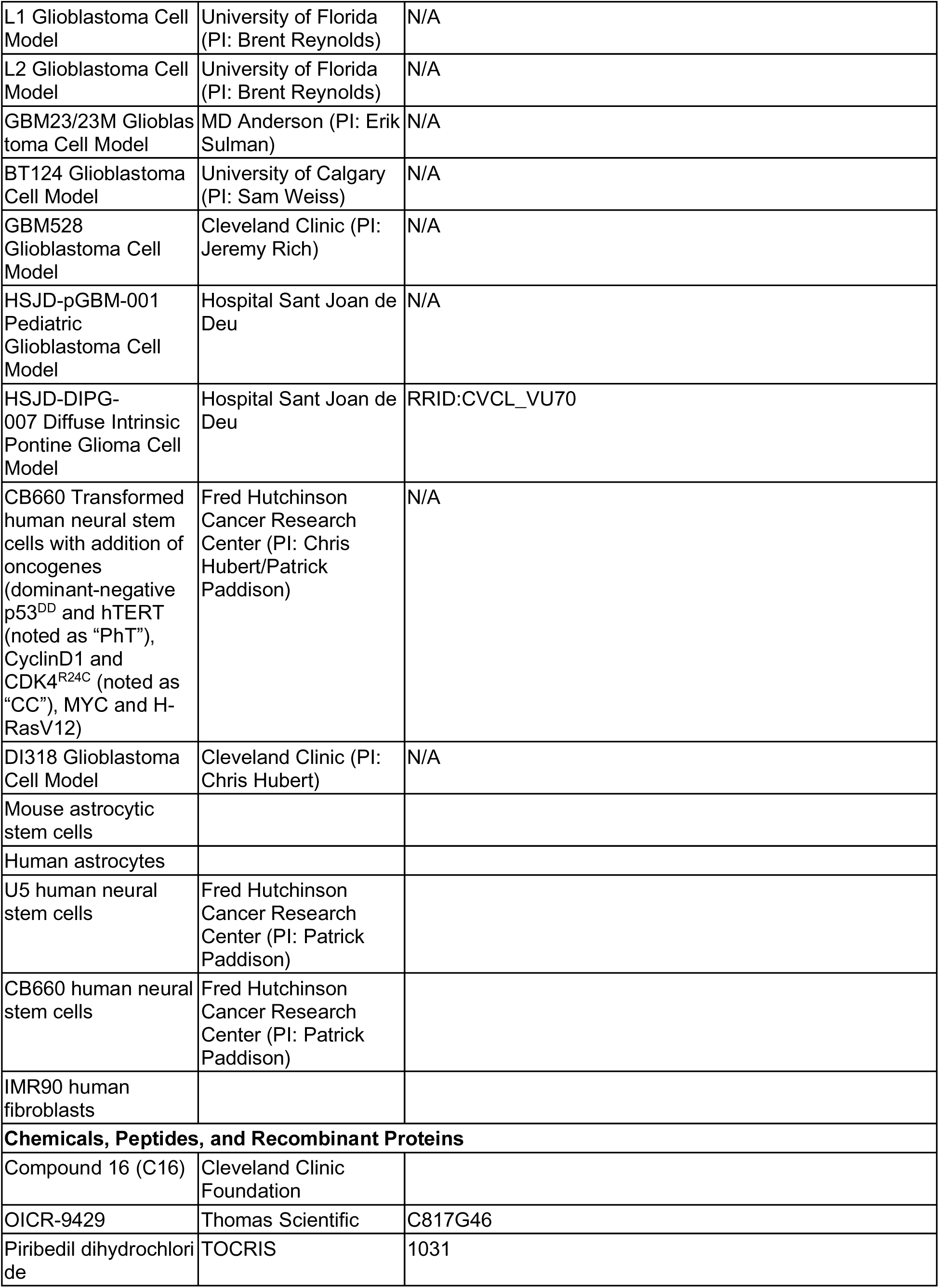
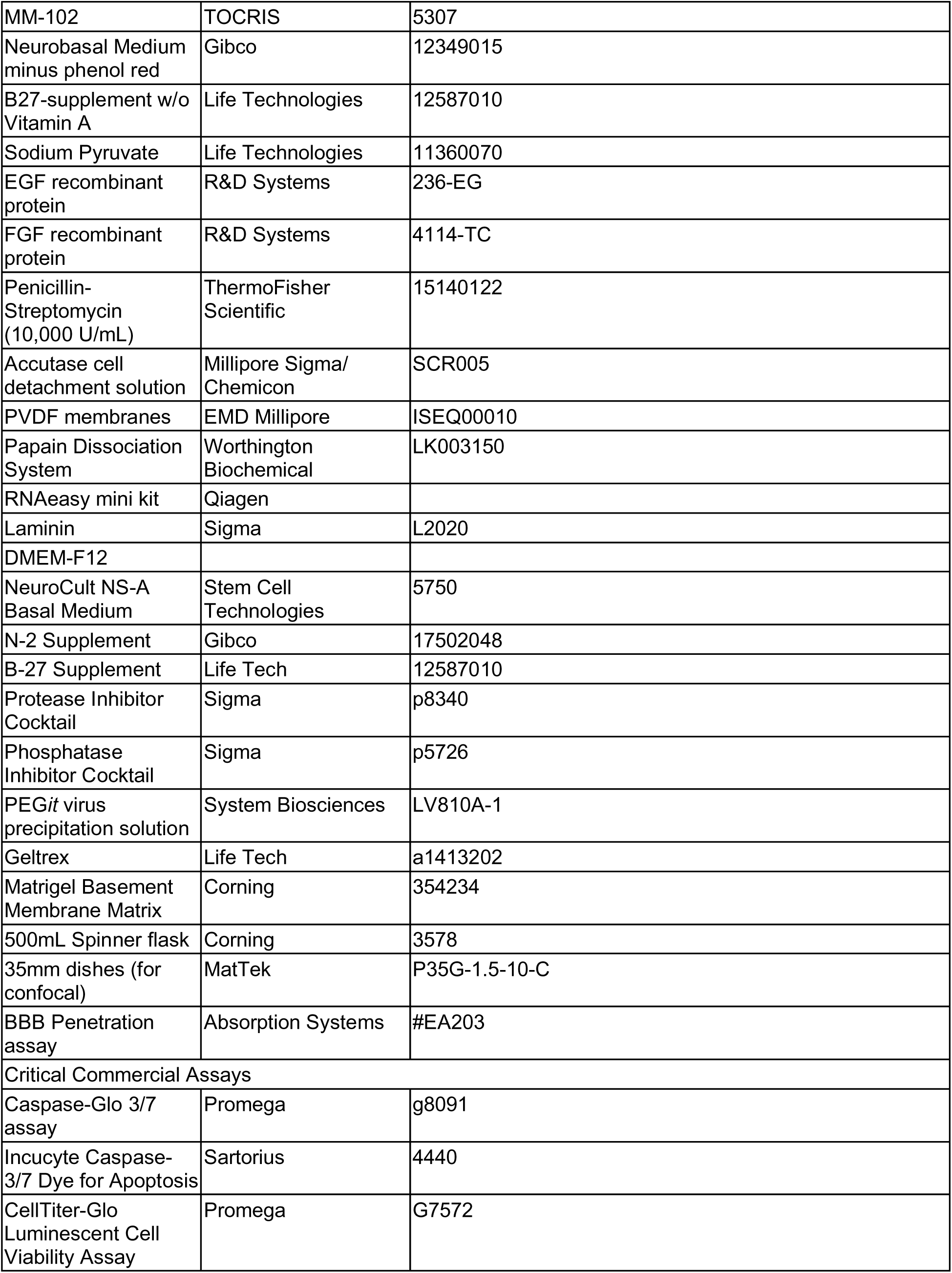
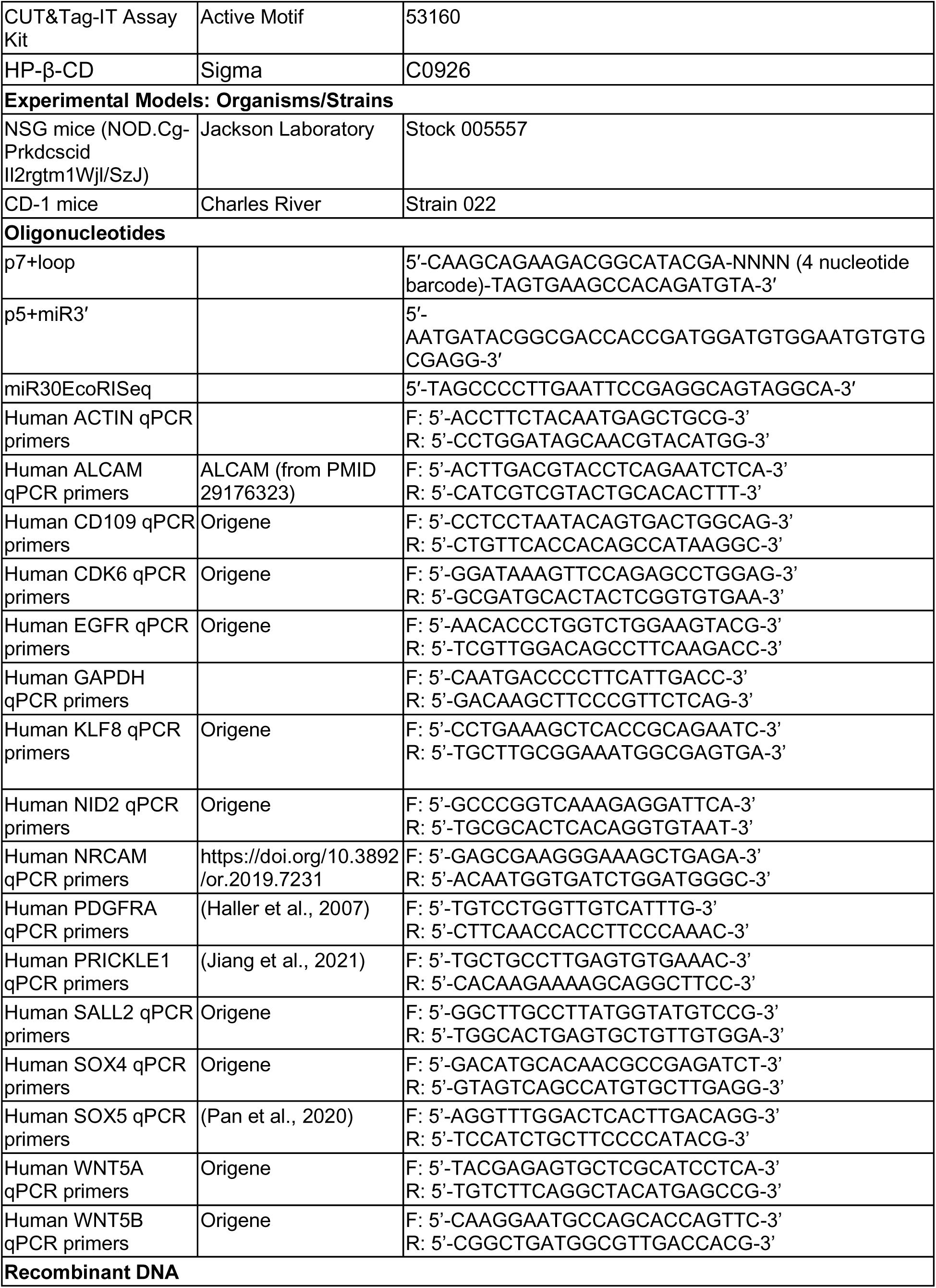
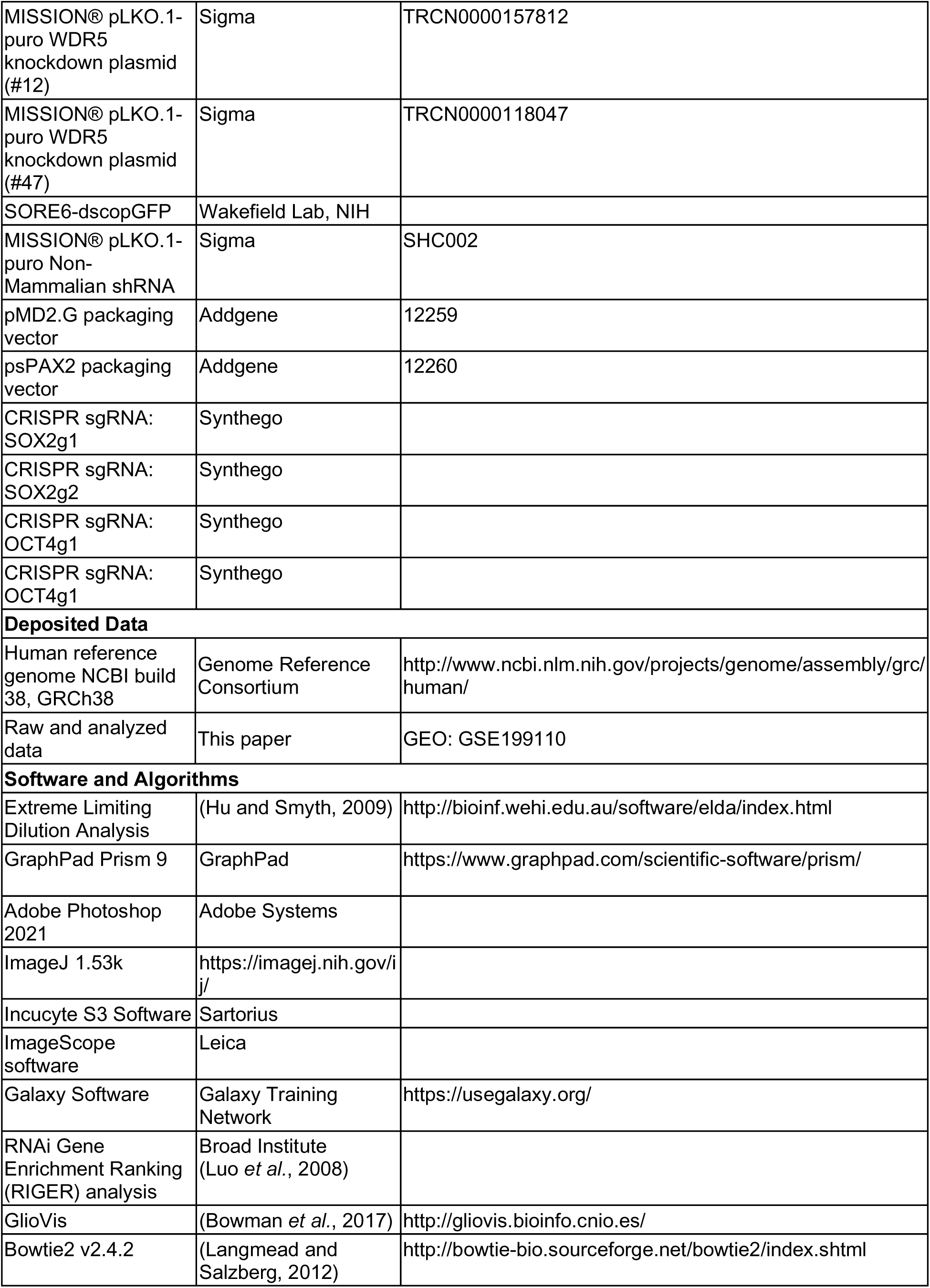
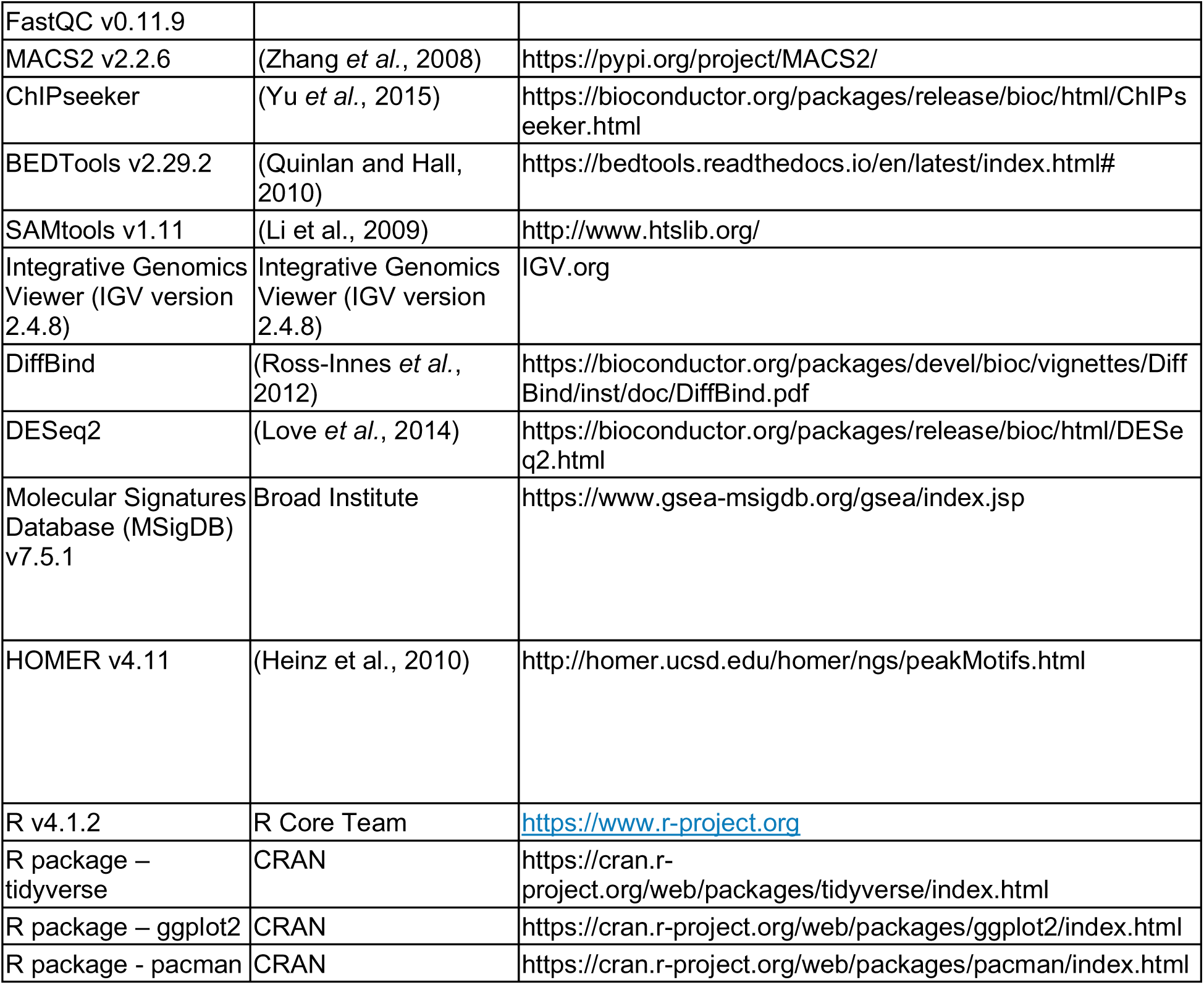

## Supplemental material

**Supp. Fig. 1** includes data in support of Figure 1 and describes the spatial expression of SOX2 I GBM organoids and validation of SORE6-GFP system in GBM CSCs. **Supp. Fig. 2** includes data in support of Figure 2 and describes the spatial functional genomics screening and hits in the SOX2-depleted niches. **Supp. Fig. 3** includes data in support of Figure 3 and describes the effect of C16 on the interaction between WDR5 and WRAD complex members. **Supp. Fig. 4** includes data in support of Figure 4 and describes H3K4me3 CUT&Tag analysis of DI318 CSCs after C16 treatment. **Supp. Fig. 5** includes data in support of Figure 5 and describes the effects of C16 on GBM CSCs expressing SOX2/OCT4. **Supp. Fig. 6** includes data in support of Figure 6 and describes expression of WDR5 in GBM patient data, the effects of WDR5 knockdown on GBM CSC self-renewal and the effects of WDR5 inhibitor MM-102 on GBM organoids and CSC culture. **Supp. Fig. 7** includes data in support of Figure 6 and describes toxicity studies and brain penetrance of C16 in mice. **Supp. Fig. 8** includes data in support of Figure 7 and describes properties of the WDR5 C16 inhibitor C16 and the effects of C16 on GBM CSC apoptosis and self-renewal and SOX2+ populations in organoids. **Supp. Table 1** includes data in support of Figure 6 and shows the list of CUT&Tag peaks unique to each treatment group (DI318 DMSO and DI318 C16). **Supp. Table 2** includes data in support of Figure 6 and shows the list of consensus CUT&Tag peaks in DI318 DMSO and DI318 C16 groups and differential enrichment analysis of these peaks between DMSO and C16 groups by DEseq2.

