## Supplemental Figures for "WDR5 represents a therapeutically exploitable target for cancer stem cells in glioblastoma"

#### Supplemental Figures and Legends

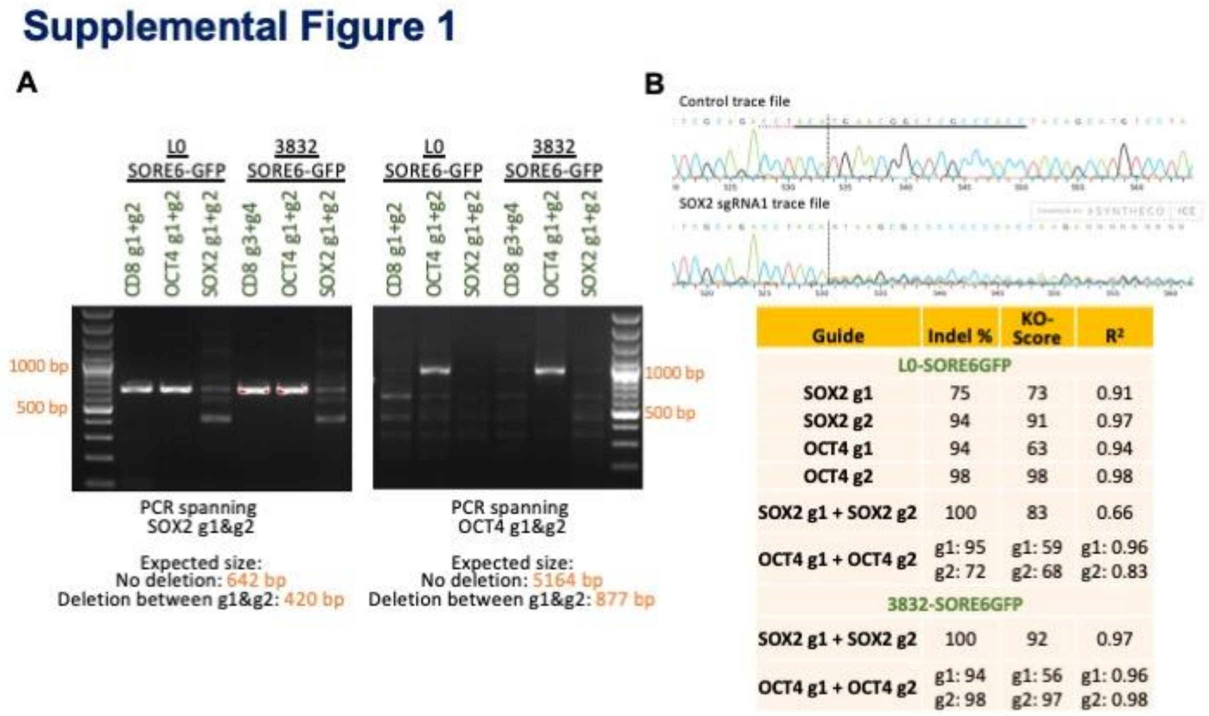


**Supplemental Figure 1 (related to Figure 1)**

(**A**) PCR amplification of genomic DNA from CSCs nucleofected with Cas9:sgRNA complexes targeting SOX2 or OCT4. Primer were designed in regions surrounding both CRISPR sgRNAs for SOX2 (left) and OCT4 (right) genes. A reduction in size indicates genomic deletion between the two sgRNA targeting sites. (**B**) Top: Example trace files from sanger sequencing of control cells and cells nucleofected with Cas9 and a SOX2 sgRNA. Guide RNA sequence is underlined and PAM sequence is dotted underlined. Bottom: Table shows indel frequencies for cells nucleofected with Cas9 and single sgRNAs or 2 sgRNAs targeting SOX2 or OCT4 (calculated using Inference of CRISPR edits analysis (Synthego ICE tool)). As SOX2 guides were close together, indel % cannot be calculated for individual guides.

#### Supplemental Figure 2

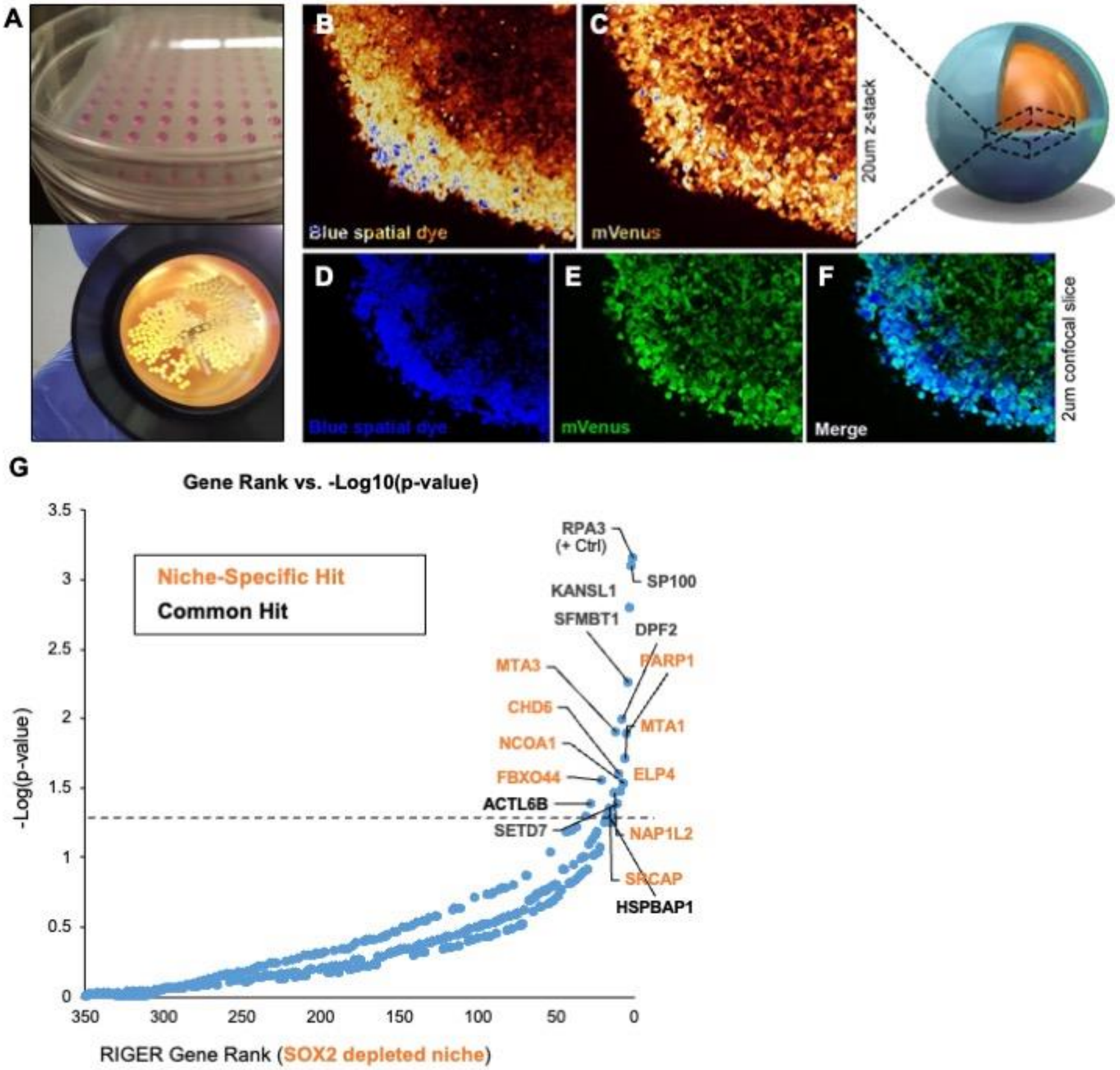

**Supplemental Figure 2 (related to Figure 2)**

(**A**) Images of GBM organoid formation and example organoid cultures in a spinning bioreactor during screen outgrowth. (**B**) Live confocal imaging of screened GBM528 organoids was used to verify proper spatial labeling prior to subsequent dissociation. Z-stacks showing labeling intensity (pseudocolored) (**B,C**) and individual image slices showing labeling overlap (**D-F**) are shown using a 20X objective. (**G**) Rank-ordered list of genes targeted in the shRNA screen, ranked by depletion of the shRNA as detected by sequencing in the SOX2-depleted niche. Dotted line represents p = 0.05 as determined by RIGER analysis of all hairpin sequences and replicates. Niche-specific hits are color coded, and common hits (including RPA3 positive control) are shown in black.

#### Supplemental Figure 3

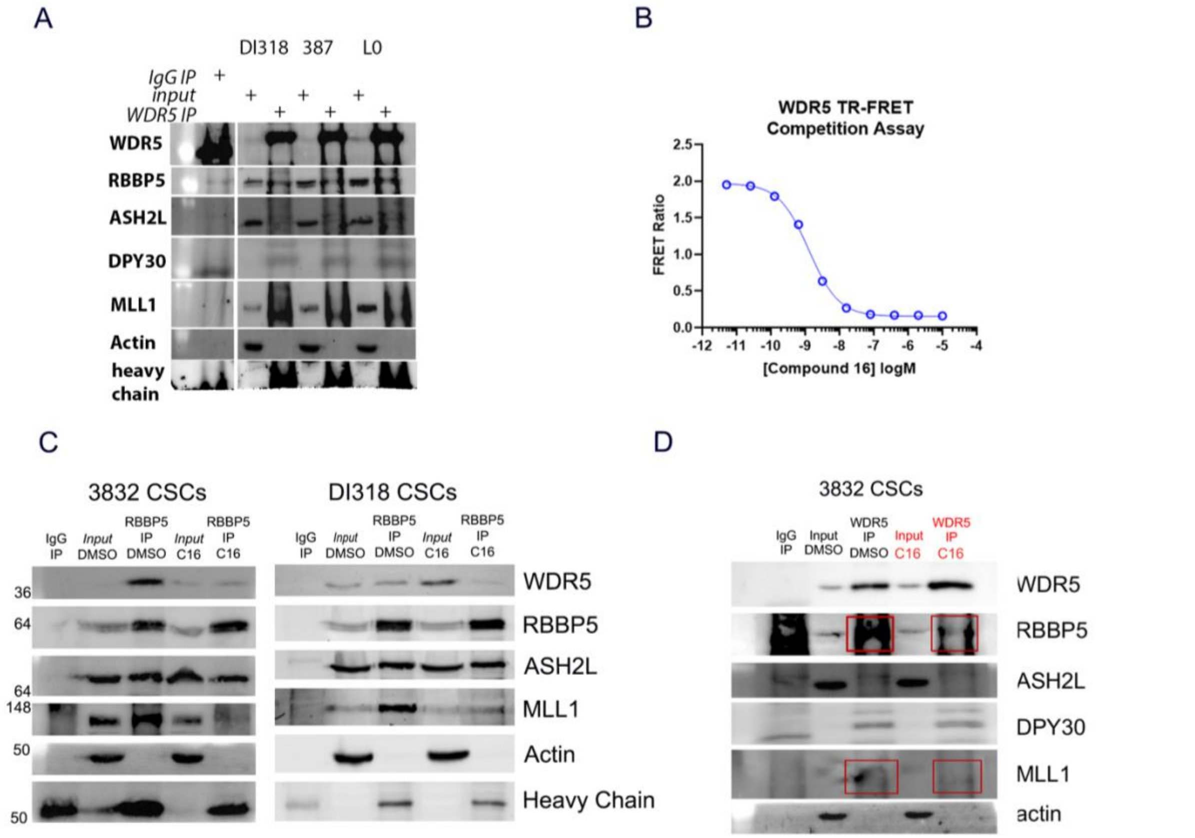


**Supplemental Figure 3 (related to Figure 3)**

(**A**) Immunoprecipitation of WDR5 in 3 GBM CSC models. Immunoblotting was performed for WRAD complex members and the WRAD-associated methyltransferase MLL1. Inputs are 5%. (**B**) Example displacement curve of compound 16 in WDR5 TR-FRET Competition Assay. Six technical replicates confirmed on-target binding with average Ki <20 pM as previously reported (Tian et al., 2020). (**C**) Additional blots related to Fig. 2C. Immunoprecipitation of RBBP5 after C16 inhibitor treatment (5 µM, 24 hrs) in 3832 and DI318 CSCs. Immunoblotting was performed for WRAD complex members and the WRAD-associated methyltransferase MLL1. Inputs are 10%. (**D**) Immunoprecipitation of WDR5 after C16 inhibitor treatment (5 µM, 1 day) in 3832 CSCs. Immunoblotting was performed for WRAD complex members and the WRAD-associated methyltransferase MLL1. Input is 5%. Representative experiments are shown. 3832 blot - input is 5%. DI318 blot – input is 10%.

### Supplemental Figure 4

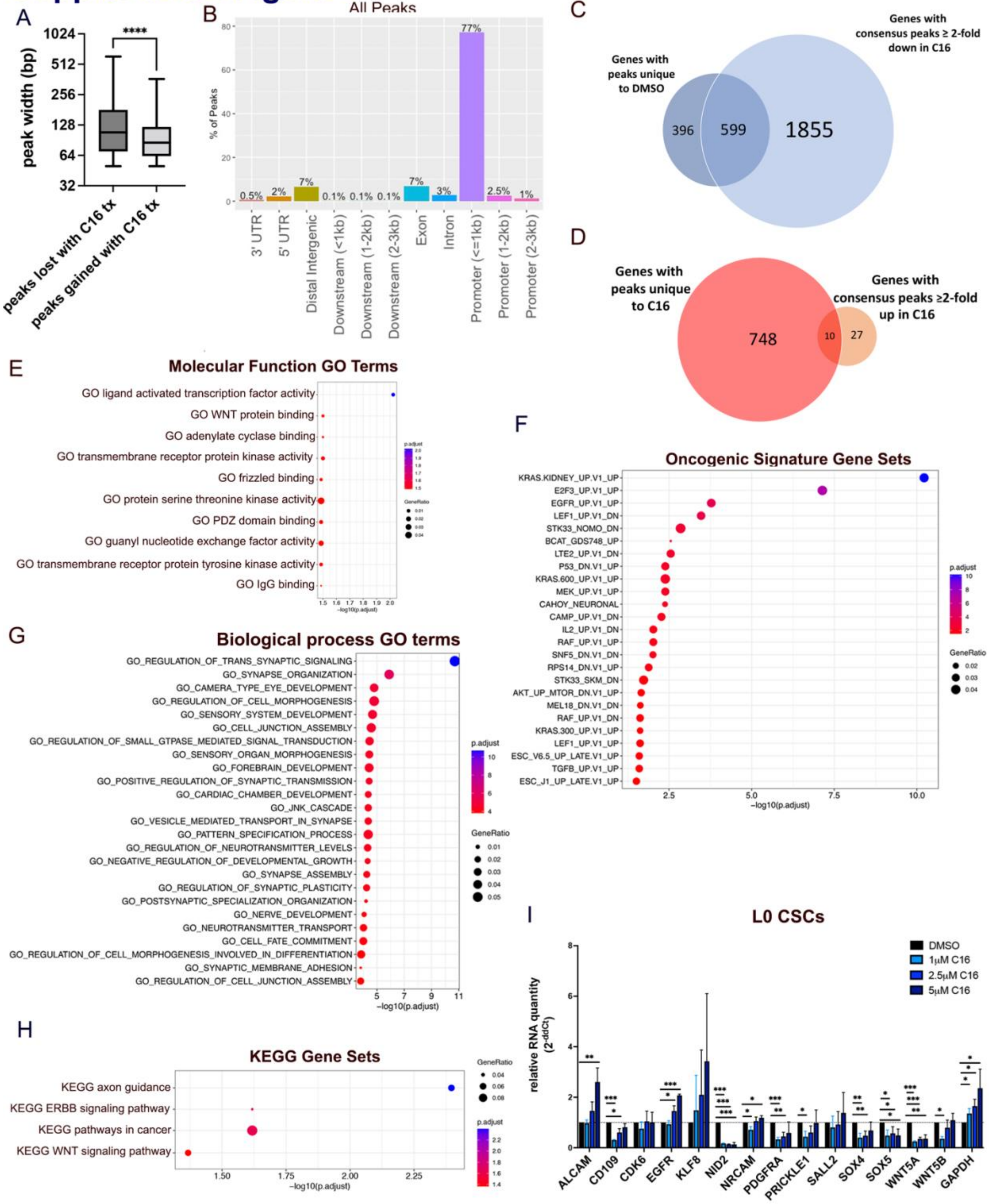

**Supplemental Figure 4 (related to Figure 4)**

**(A)** Box and whisker plots comparing peak width for CUT&Tag peaks lost in C16 treatment group (unique to DMSO) and peaks gained in the C16 treatment group (unique to C16). P value determined by KS test. **(B)** Bar plot showing distribution of all consensus peaks (from DESeq2) in gene regions. **(C-D)** Venn diagrams showing overlap between genes with unique peaks and differential peaks in each treatment group. (**E-H**) MSigDB gene set annotations enriched among CUT&Tag peaks decreased (log2FC≤-1) after C16 treatment of DI318 cells. (**I**) qPCR for specified genes on L0 CSCs treated with indicated doses of C16 for 72 hrs. Bars represent mean expression of n=3 biological replicates, normalized to ACTB levels by ddCt method, +/- SD.

### Supplemental Figure 5

A

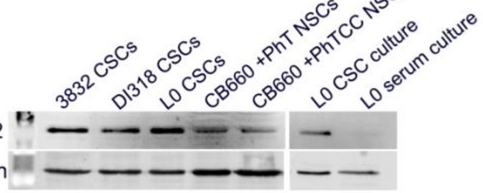


B

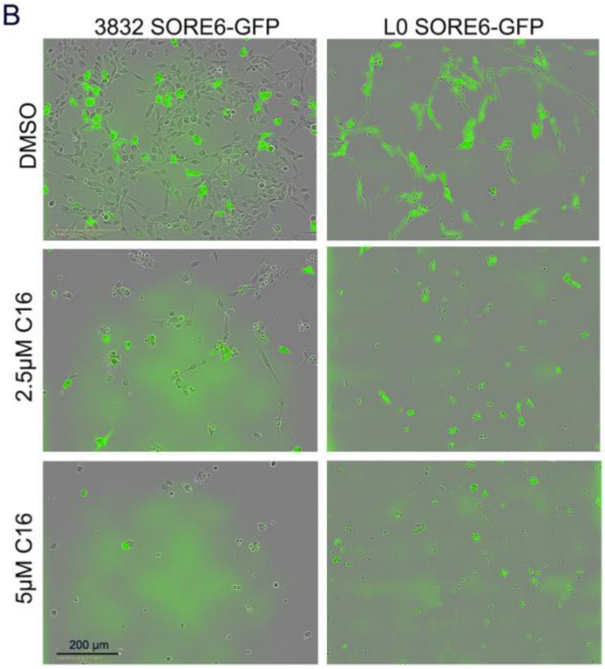


C

| Cell model | Average sphere forming frequency | |
|---|---|---|
| | *GFP low* | *GFP high* |
| DI318-SORE6GFP (n=5) | 1 in 4.4 | 1 in 2.4 |
| L0-SORE6GFP (n=5) | 1 in 3.9 | 1 in 2.0 |

D

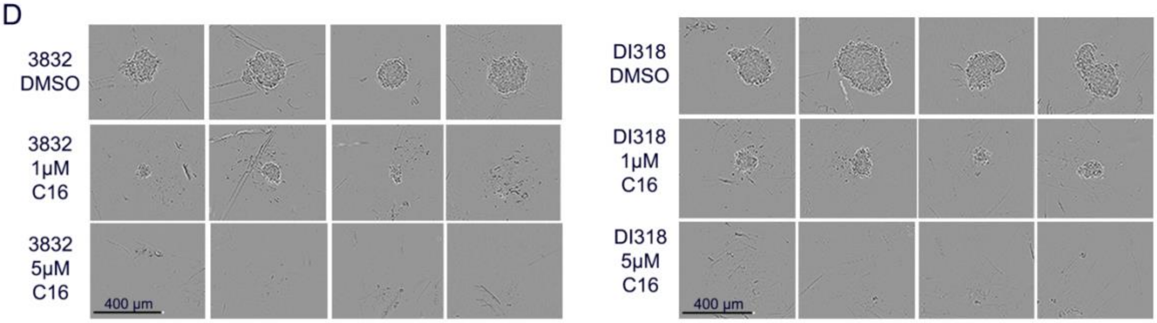


E

| Cell model | Average sphere forming frequency | | |
|---|---|---|---|
| | *DMSO* | *1uM C16* | *5uM C16* |
| **DI318** (n=5) | 1 in 1.9 | 1 in 5.6 | 1 in 50.0 |
| **3832** (n=4) | 1 in 2.3 | 1 in 4.2 | 1 in 33.3 |
| **L0** (n=4) | 1 in 1.4 | 1 in 33.3 | 1 in 333.3 |

F

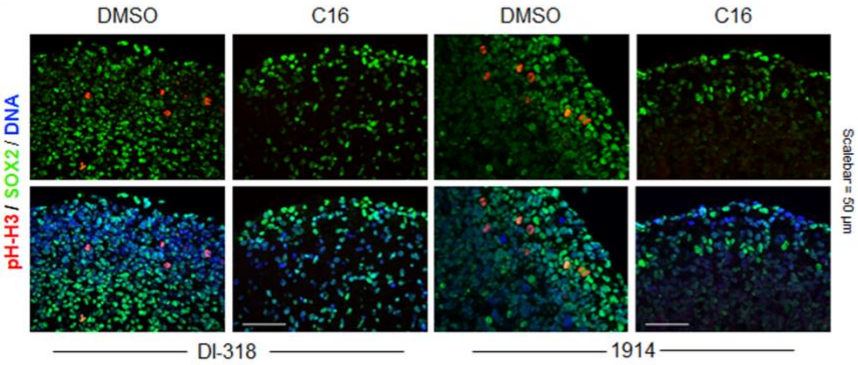

**Supplemental Figure 5 (related to Figure 5)**

(**A**) SOX2 expression in 3 CSC models, transformed human neural stem cells (PhT, PhTCC, refer to Figure 6C and associated text for details), and L0 CSCs grown in CSC-enriching culture conditions (“CSC culture”) and differentiation (DMEM+10% FBS) conditions (“serum culture”). (**B**) GBM CSC models transduced with the SORE6-GFP reporter were treated with the C16 WDR5 inhibitor. Representative images Day 7 post treatment are shown. (**C**) *In vitro* limiting-dilution analysis was performed on $GFP^{low}$ and $GFP^{high}$ populations isolated by FACS from bulk SORE6-GFP-transduced cells. Sphere-formation frequency from multiple independent replicates is shown. (**D-E**) *In vitro* limiting-dilution analysis was performed on CSCs in the presence of C16. (**D**) Representative spheres from limiting-dilution assays. (**E**) Sphere-formation frequency from multiple independent replicates is shown. (**F**) Immunostaining of C16-treated GBM organoids (10 µM for 7 days) for SOX2 and pHH3 to identify proliferative SOX2+ cells.

### Supplemental Figure 6

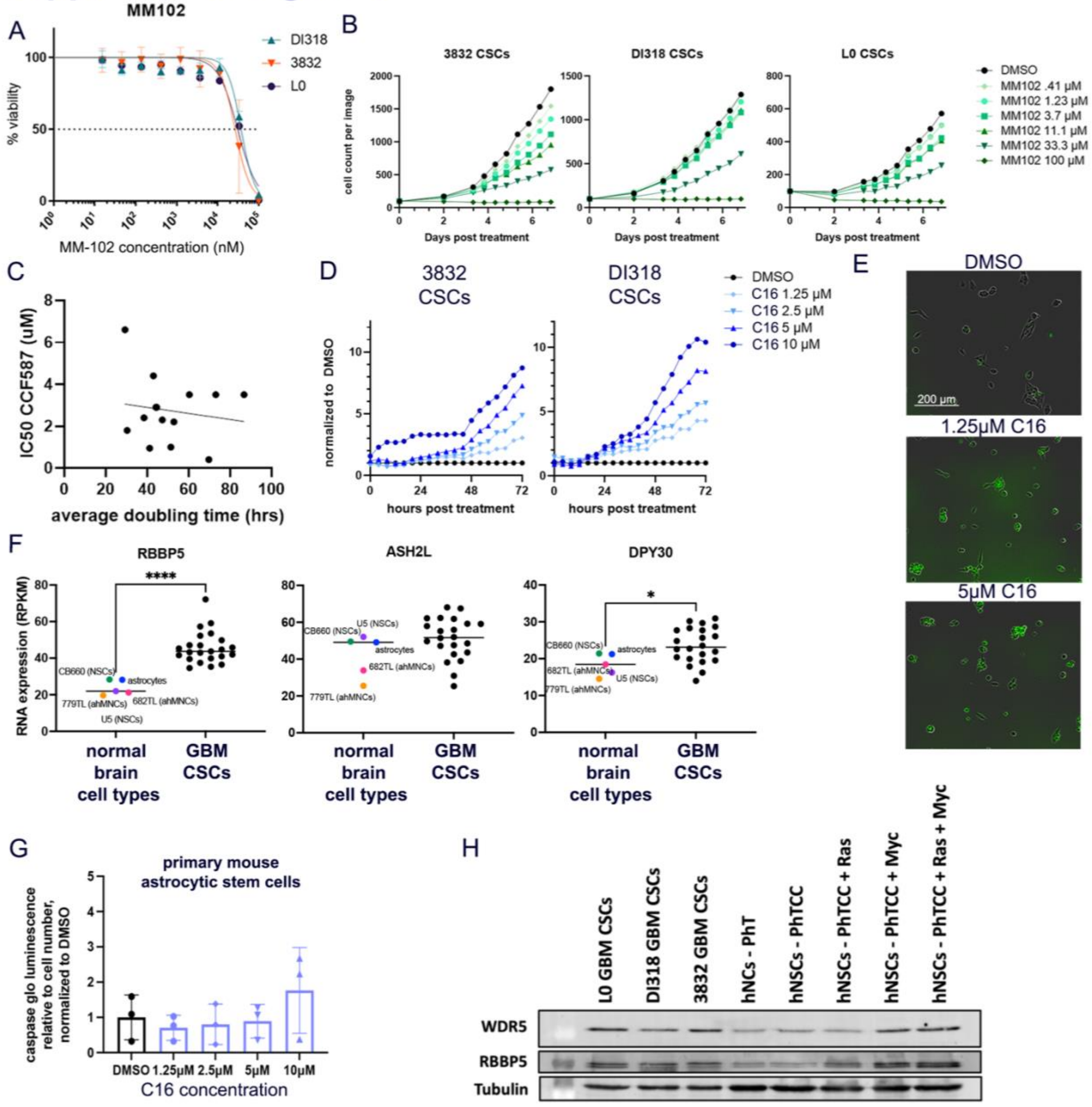


**Supplemental Figure 6 (related to Figure 6)**

(**A**) CSCs were treated with a range of concentrations of MM-102, a peptide inhibitor of WDR5. After 7 days, viable cell counts were measured by CellTiter Glo viability assay. Values represent mean luminescence values normalized to DMSO-treated cells. (**B**) Proliferation of MM-102-treated CSCs over 7 days, determined by IncuCyte live cell imaging. Values represent mean cell count per image, +/- SD; n=3 technical replicates; one representative experiment is shown per

CSC model. (**C**) Doubling times and average IC50s of C16 are plotted for multiple CSC models and transformed neural stem cell models. Line of best fit (simple linear regression) is shown. $R^2$=0.022. (**D-E**) CSCs were treated with a range of concentrations of C16, and Caspase-3/7 green dye reagent (Sartorius) was added to culture medium. (**D**) Representative images 4 days after treatment with C16 are shown. (**E**) Quantification of green cells relative to % confluence for each dose is graphed (normalized to DMSO treatment group) over 3 days, determined by IncuCyte live cell imaging. One representative experiment is shown per CSC model. (**F**) WRAD complex member expression from RNA sequencing of a panel of normal brain cell types and GBM lines (from (Toledo et al., 2015)). Each point represents average expression from multiple sequencing replicates. (**G**) Primary mouse astrocytic stem cells were treated with a range of concentrations of C16 and subjected to Caspase 3/7 Glo luminescence assay after 4 days to measure caspase 3/7 activity. Bars represent fold change in caspase 3/7 activity per cell relative to the average for DMSO-treated cells, +/- SD; circles represent biological replicates. (**H**) WDR5 and RBBP5 member expression in 3 CSC models and transformed human neural stem cells (hNSCs).

#### Supplemental Figure 7

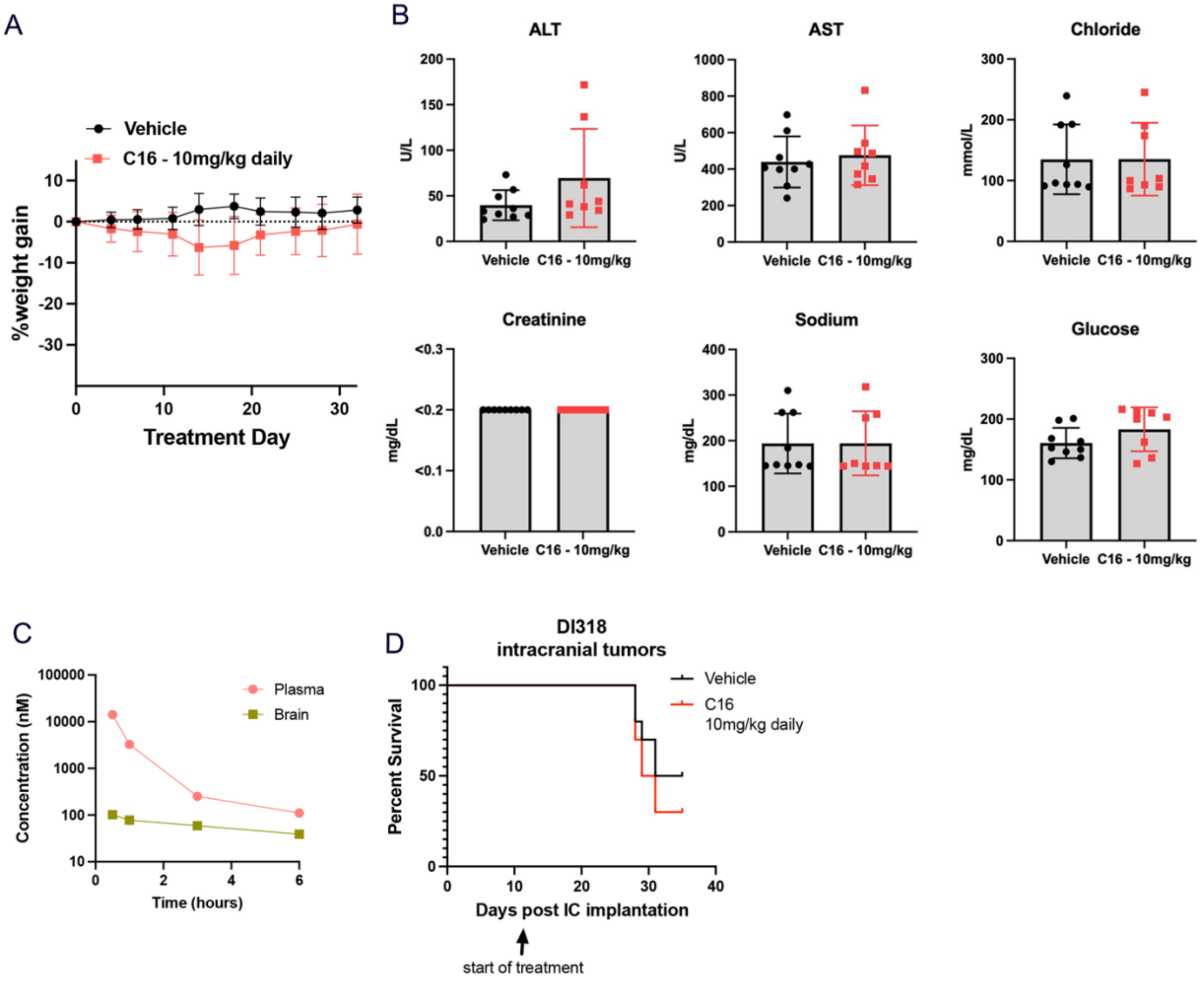


**Supplemental Figure 7 (related to Figure 6)**

(**A**) C16 (10 mg/kg) was injected intraperitoneally in NSG mice every day for 30 days, and mice were weighed twice weekly. (**B**) Blood chemistry panels on mice treated for 30 days with 10 mg/kg C16 (from (A)). (**C**) C16 (10 mg/kg) was injected intraperitoneally in CD-1 mice, brains/plasma were collected at various time points, and the quantity of C16 in each tissue was measured by mass spectrometry. (**D**) Kaplan-Meier survival plot of mice intracranially implanted with DI318 CSCs and treated with C16 daily. n=10 per group.

### Supplemental Figure 8

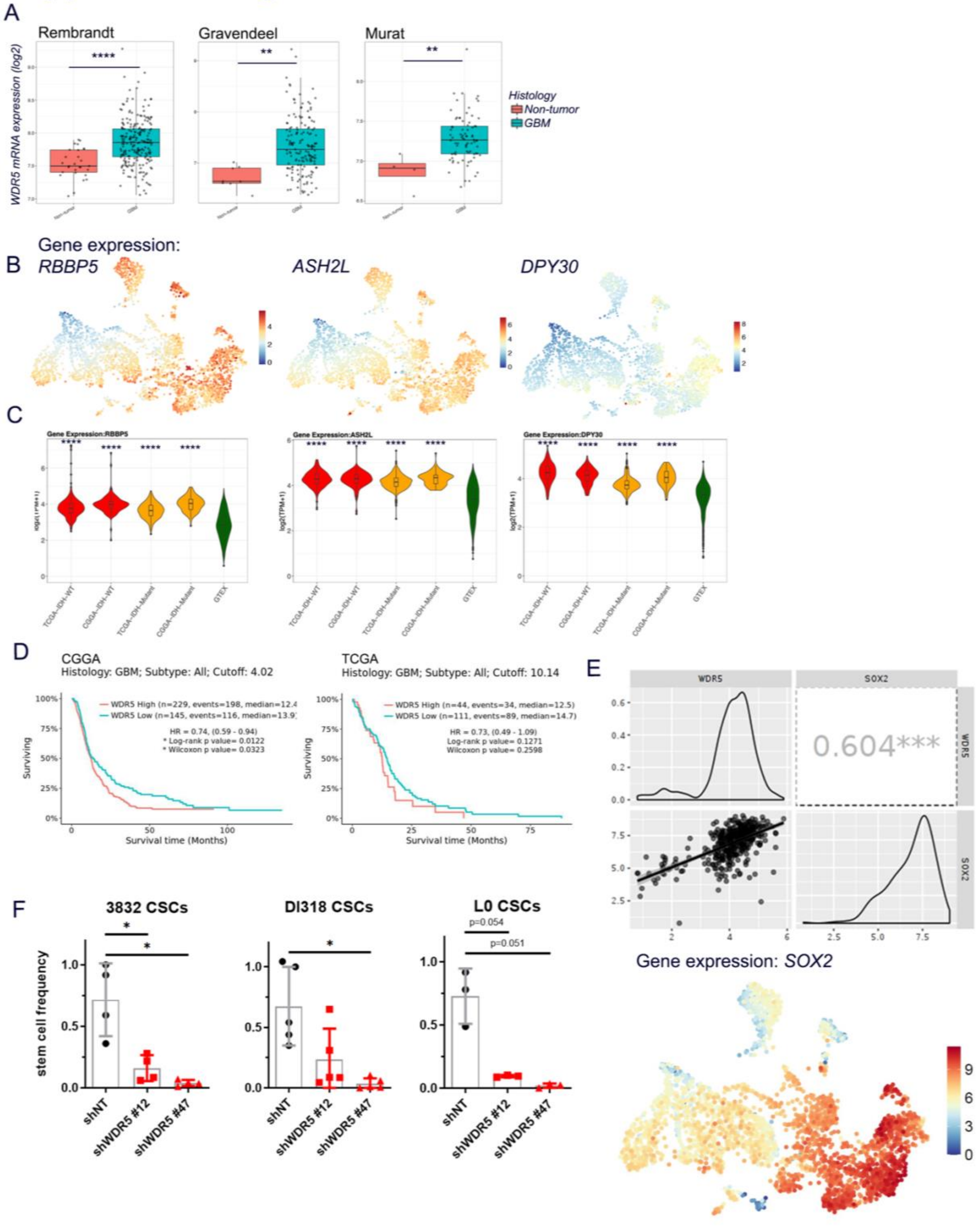


**Supplemental Figure 8 (related to Figure 7)**

(**A**) WDR5 RNA expression in GBM and normal brain specimens from various RNA expression datasets. Pairwise comparisons (unpaired t-test) were performed between groups with Bonferroni corrections for multiple testing. P-values; Rembrant: p=2.0E-06; Gravendeel: p=1.4E-03; Murat: p=6.2E-03. Data obtained from the GlioVis database. (**B**) UMAP projections of gene expression for WRAD complex members on BRAIN-UMAP. (**C**) RNA expression of WRAD complex members in selected tumor and normal groups. p values determined by unpaired t-tests with correction for multiple comparisons. (**D**) Overall survival of GBM patients graphed based on WDR5 expression from Chinese Glioma Genome Atlas (CGGA) and The Cancer Genome Atlas (TCGA) datasets. Accessed via GlioVis database. (**E**) Top: Co-expression of WDR5 and SOX2 in GBM from the Chinese Glioma Genome Atlas (n=225 wild-type IDH1 GBM samples, accessed via GlioVis database). Scatter plots (lower left), density plots (middle diagonal), Pearson's correlation coefficients with statistical significance (***p<0.001; upper right) are provided on the plot. Bottom: UMAP projection of *SOX2* gene expression on BRAIN-UMAP. (**F**) *In vitro* limiting-dilution analysis was performed on WDR5 KD cells and shNT control cells. Bars represent mean sphere-formation frequency, +/- SD; symbols represent biological replicates.

**Supplemental Table 1**

CUT&Tag peaks ≥50 bp unique to each treatment group (DI318 DMSO and DI318 C16). Data was generated with BedTools.

**Supplemental Table 2**

List of consensus CUT&Tag peaks in DI318 DMSO and DI318 C16 groups and differential enrichment analysis of these peaks between DMSO and C16 groups by DEseq2. **Sheets**: Sheet 1: shows all peaks; Sheet 2: shows only significant peaks (FDR < 0.05); Sheet 3: shows only significant peaks in C16 group (FDR < 0.05 & fold > 0); Sheet 4: shows only significant peaks in DMSO group (FDR < 0.05 & fold < 0). **Column headings**: Log2Conc: mean read concentration over all the samples (log2 normalized CUT&Tag read counts); log2(conc_C16): mean concentration over the C16 treatment group; log2(conc_DMSO): mean concentration over the DMSO treatment group; log2(ratio conc_C16/conc_DMSO): shows the difference (log2FC) in mean concentrations between the two treatment groups, with a positive value indicating enrichment in the C16 treatment group and a negative value indicating enrichment in the DMSO treatment group. Data was generated by DiffBind and DESeq2.
